# Unraveling Lineage Roadmaps and Cell Fate Determinants to Postnatal Neural Stem Cells and Ependymal cells in the Developing Ventricular Zone

**DOI:** 10.1101/2024.06.16.599182

**Authors:** Jianqun Zheng, Yawen Chen, Yukun Hu, Jie Lin, Yujian Zhu, Manlin Xu, Weihong Song, Xi Chen

## Abstract

The ventricular zone (VZ) is made up of adult neural stem cells (NSCs) and multi-ciliated ependymal cells (EPCs). Both NSCs and EPCs are derived from radial glial cells (RGCs). To date, the transcriptomic dynamics and the molecular mechanisms guiding the cell fate commitment during the differentiation remain poorly understood. In this study, we analysed the developing VZ at the single-cell resolution and identified three distinct cellular states of RGCs: bipotent glial progenitor cells (bGPCs), neonatal NSC-neuroblasts (nNSC-NBs) and neonatal EPCs (nEPCs). The differentiation from bGPCs to nNSC-NBs and nEPCs forms a continuous bifurcating trajectory. Further molecular analysis along the NSC branch unveiled a novel intermediate state of cells expressing oligodendrocyte precursor cell (OPC) and neuroblast (NB) marker genes. Several transcription factors (TFs) were proved to be essential for the EPC-lineage differentiation, with TFEB emerging as a key regulator dictating the bGPC fate. The activation of TFEB promoted differentiation towards NSC-NBs while restrained the trajectory towards EPCs. Our findings offer detailed insights into further understanding VZ development and lay the groundwork for investigating potential therapeutic strategies against VZ-related disorders, such as hydrocephalus and neurodegenerative diseases (NDDs).

## Introduction

The ventricular system, consisting of inter-connected cavities filled with cerebrospinal fluid (CSF), is a unique characteristic of the brain in vertebrates [1]. The cell layers lining the ventricle are referred to as the ventricular zone (VZ), which is the primary region for neurogenesis and gliogenesis in mammals [2, 3]. Mice have a similar ventricular system to humans where four major cavities are observed in both species, making mice an ideal model for the functional and developmental study of the ventricular system.

During the mid-to-late stages of mouse embryonic development, the VZ is predominantly occupied by radial glial cells (RGCs) that are named after their common radial appearance [2]. High heterogeneity was observed during the development of RGCs in the VZ where the majority of RGCs participated in neurogenesis and only one sixth of them contributed to gliogenesis [4]. Around embryonic day 15, the glial-fated RGCs undergo their final mitosis in the embryonic stage, giving rise to glial progenitor cells (GPCs). These GPCs maintain their radial glial morphology until birth [5, 6]. In the neonatal stage, some GPCs migrate to the cortex and differentiate into oligodendrocytes and astrocytes, while others remain in the VZ and differentiate into neural stem cells (NSCs) and ependymal cells (EPCs) [7].

NSCs have stem cell characteristics and are capable of self-renewal and differentiation into mature neurons. During the differentiation process, the intermediate transitional states of transit amplifying cells and neuroblasts (NBs) can be observed [8–11]. These regenerating features make NSCs with great potential values in treating neurodegenerative diseases (NDDs), brain injuries, and strokes [12]. In the VZ of the adult brain in mammals, NSCs are surrounded by EPCs that have multiple motile cilia. As the apical domain of EPCs is much larger than that of NSCs, the VZ manifests a unique pinwheel organisation [13]. Cilia are microtubule-based organelles protruding from the cell surface [14]. The rhythmic beating of cilia of EPCs facilitates the circulation of CSF, which is critical for supplying nutrients and removing metabolic waste from brain cells and hence has a great impact on the metabolism, the self-renewal and the differentiation of NSCs [15–18]. Disruption of the circulation of CSF often leads to hydrocephalus [19].

Although the VZ represents the largest germinal niche in the adult mammalian brain [2, 3], the postnatal developmental process of the VZ remains poorly understood. Recent studies with advanced lineage-tracing methods have provided valuable insights on the clonal relationships among cells in the VZ during the development [6, 20–22]. However, a holistic picture of the transcriptomic dynamics during the development of the VZ is still lacking, and mechanisms underlying the cell fate commitment during the differentiation remain elusive. Here we performed single-cell RNA-sequencing (scRNA-seq) and transcription factor (TF) ChIP-seq to depict the detailed molecule events along the bifurcating differentiation trajectories from GPCs to NSCs and EPCs. Several TFs and pathogenic genes emerged involved in determining the fate of VZ progenitors. Of note, we discovered that TFEB, a master regulator of lysosome biogenesis [23, 24], plays pivotal roles in cell fate specification within the VZ, suggesting a novel treatment strategy of targeting TFEB during development for VZ-related disorders such as NDDs.

## Results

### Single-cell transcriptomic data unveil the bifurcating differentiation trajectories of RGCs in the postnatal VZ

To reveal the cellular heterogeneity of RGCs and their differentiation roadmaps to postnatal NSCs and EPCs, scRNA-seq techniques were applied on the developing VZ from mice. The neonatal stage was selected for three practical reasons. First, RGCs stop producing neurons after birth [25], eliminating the influence of neurogenesis. Second, cells responsible for generating oligodendrocytes and astrocytes have migrated to the cortex during this stage [7, 20]. Finally, lineage tracing experiments have demonstrated that RGCs from the neonatal stage possess the bipotency to differentiate into NSCs and EPCs [21, 22].

CD133 has been shown to be specifically expressed on the apical surface of RGCs in the neonatal mouse VZ [26–28], which serves as an excellent marker for RGC isolation. Therefore, VZ-containing brain tissues at the neonatal stage were dissected under a stereoscope and single cell suspension was prepared and stained with an anti-CD133 antibody (see **Methods**). CD133-positive RGCs were isolated using flow cytometry sorting (FACS). Subsequently, scRNA-seq libraries were prepared using the 10X Genomics (10X) platform (**Figure 1A** and **Supplementary Figure 1A**). In addition, small-scale experiments were also performed independently to confirm the findings using a plate-based 3’ scRNA-seq method that was developed previously in our lab [29].

**Figure 1.**
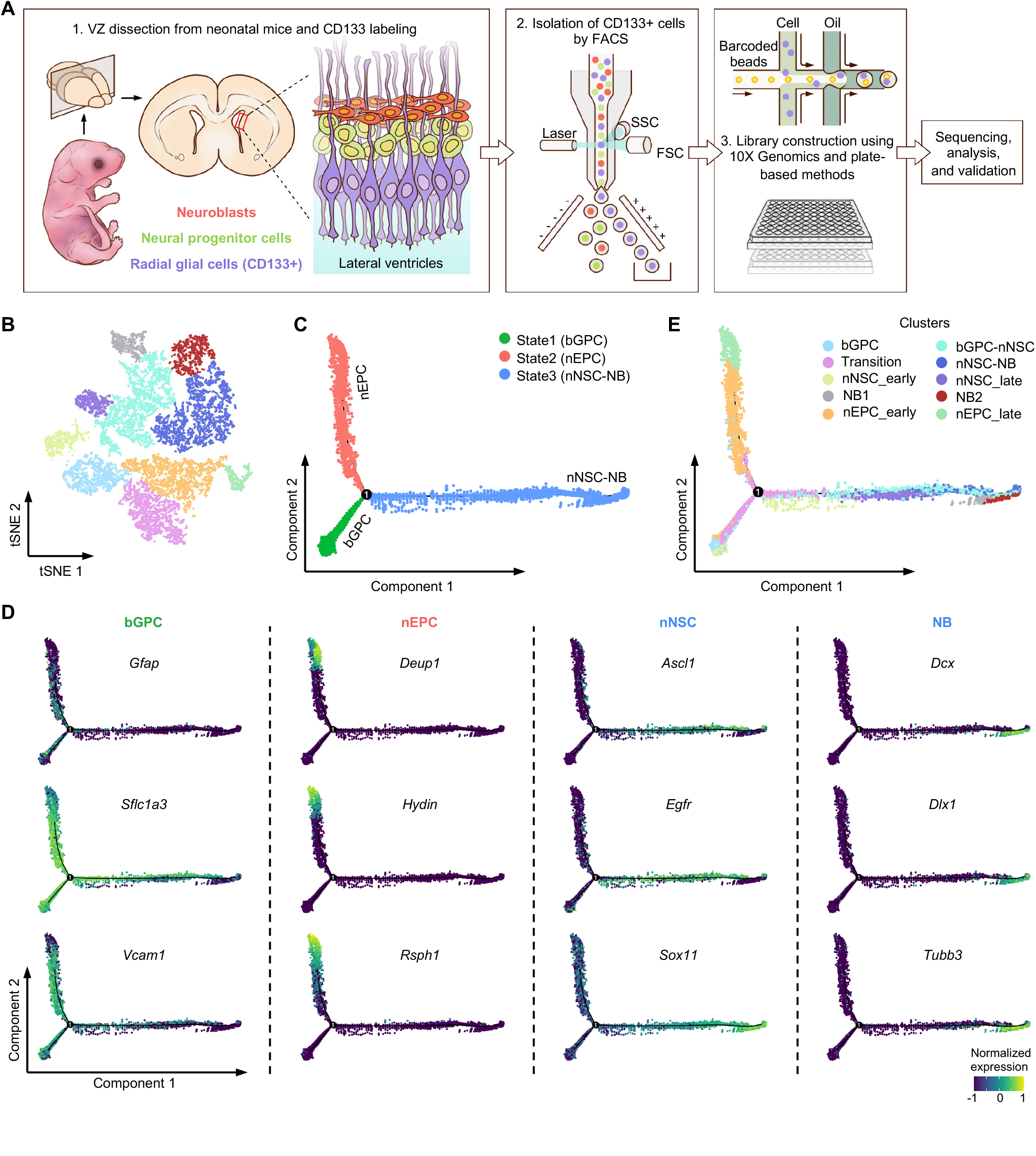
The developmental trajectory of postnatal ventricular zone by single-cell transcriptomics. **(A)** Schematic diagram of the experimental workflow. Radial glial cells (RGCs), labeled by CD133, were isolated from the ventricular zone (VZ) of neonatal mice by fluorescence-activated cell sorting (FACS). Single-cell RNA sequencing (scRNA-seq) libraries were then constructed using droplet-based 10X Genomics (10X) and plate-based modified Smart-seq3 (Plate) method, followed by sequencing, analysis and validation. SSC, side scatter; FSC, forward scatter. **(B)** t-SNE projection of 13,223 cells in the 10X data showing the transcriptional similarities among RGCs at the neonatal stage. **(C)** Pseudotime analysis of all RGCs at the neonatal stage showing a bifurcating differentiation trajectory of the VZ development. Different states are coluor-coded. The solid black circle indicates the branching point. bGPC, bipotent glial progenitor cell; nEPC, neonatal ependymal cell; nNSC, neonatal neural stem cell; NB, neuroblast. **(D)** Expression profiles of representative markers along the bifurcating trajectory. **(E)** The same bifurcating differentiation trajectory as shown in **(C)**. Cells are coloured by the cluster shown in **(B)**.

Overall, a total of 13,223 and 2,594 cells were obtained from 10X and the plate-based methods after quality control (see **Methods**), respectively. The data qualities of the two methods were similar (**Supplementary Figure 1B**). The Unique Molecular Identifier (UMI) count per cell exceeded 10,000, and the numbers of detected genes number ranged from 2,000 to 6,000 (**Supplementary Figure 1B**). Unsupervised clustering analysis on the 10X data identified 10 distinct clusters, representing different subtypes of GPCs, NSCs, EPCs and some transitional cell states (**Figure 1B**). The same clusters were also observed in the plate-based data (**Supplementary Figure 1C**), indicating the robustness of the results. To investigate the relationship among the cells during the development of VZ, a pseudotime analysis was performed using Monocle [30] on the 10X data, and a bifurcating differentiation trajectory was clearly observed (**Figure 1C**). A very similar pseudotime trajectory with the same bifurcation was also observed (**Supplementary Figure 1D**) using the diffusion map algorithm implemented in destiny [31]. In addition, the same results could also be observed in the independently performed plate-based data (**Supplementary Figure 1C**), which serves as a validation for the differentiation trajectory. Due to the larger number of cells, we mainly focused on analysing the 10X data subsequently.

Three main states were observed along the differentiation trajectory (**Figure 1C**). Many gliogenic marker genes including *Gfap*, *Slc1a3*, *Vcam1*, *Aldh1l1*, *Aqp4* and *Fabp7* were present in state 1 cells, with the highest expressions observed in cells at the bottom-left corner of the 2D plot (**Figure 1C**, **D** and **Supplementary Figure 1E**). Therefore, it was likely state 1 cells were GPCs and they were the “root” of the differentiation trajectory. High and specific expressions of well-known EPC markers like *Deup1*, *Hydin*, *Rsph1*, *Dynlrb2*, *Ift20*, and *Tekt4* were detected in state 2 cells (**Figure 1D** and **Supplementary Figure 1E**). NSC markers, such as *Ascl1*, *Egfr* and *Sox11*, showed the highest expression in cells around the middle of the state 3 branch (**Figure 1D**), and neuroblast (NB) marker genes, including *Dcx*, *Dlx1* and *Tubb3*, were specifically expressed at the end of the state 3 branch (**Figure 1D**). Previous studies demonstrated that *Gmnc* and *Gmnn* are antagonistic Geminin family members. *Gmnc* expression favoured the generation of EPCs while *Gmnn* induced an NSC fate [21, 32]. Consistently, *Gmnc* and *Gmnn* showed higher expression in state 2 and state 3 branches, respectively (**Supplementary Figure 1E**). Based on the expression patterns of those genes, state 2 and state 3 branches were likely to represent the EPC and NSC-NB lineages, respectively.

Next, the 10 cell clusters identified from the clustering analysis were visualised on the pseudotime trajectory (**Figure 1E**). By looking at the marker genes and the relative positions on the pesudotime, we successfully assigned the identity of each cluster (**Figure 1E**). The whole pseudotime trajectory described the bifurcating differentiation process from bipontent GPCs (bGPCs) to neonatal EPCs (nEPCs) and neonatal NSCs (nNSCs) and NBs, allowing a detailed analysis of the expression dynamics of various cell fate marker genes along the branches (**Supplementary Figure 1F**).

### Molecular cascades underlying bGPC commitment to nNSC-NB

Next, a detailed analysis of the nNSC-NB branch was conducted to identify the genes essential for this lineage. Due to the lack of prior information on the molecular characteristics of nNSCs in the VZ, a robust set of specific markers from adult NSCs was used [8–11]. Adult NSCs include quiescent NSCs (qNSCs) that are in a resting state and activated NSCs (aNSCs) which are in a proliferative state capable of self-renewal and generating neurons under certain conditions [9]. A vast majority of qNSC markers showed higher expression levels in bGPCs and gradually declined during the bGPC to nNSC-NB branch (**Figure 2A**, top panel). In contrast, aNSC markers showed the opposite trend, where their expressions were low in bGPCs and became progressively higher during the differentiation (**Figure 2A**, bottom panel). Thus, the conversion from bGPCs into nNSCs was reminiscent of the transition from qNSCs to aNSCs in adulthood. Previous study showed that the VZ growth during juvenile development can be explained by the increasing size of the apical domains of differentiating EPCs, despite a net loss in postnatal NSC number [22]. It can be extrapolated that bGPCs develop into qNSCs in adult stage, while nNSCs become adult aNSCs and gradually migrate out of the VZ (**Figure 2B**).

**Figure 2.**
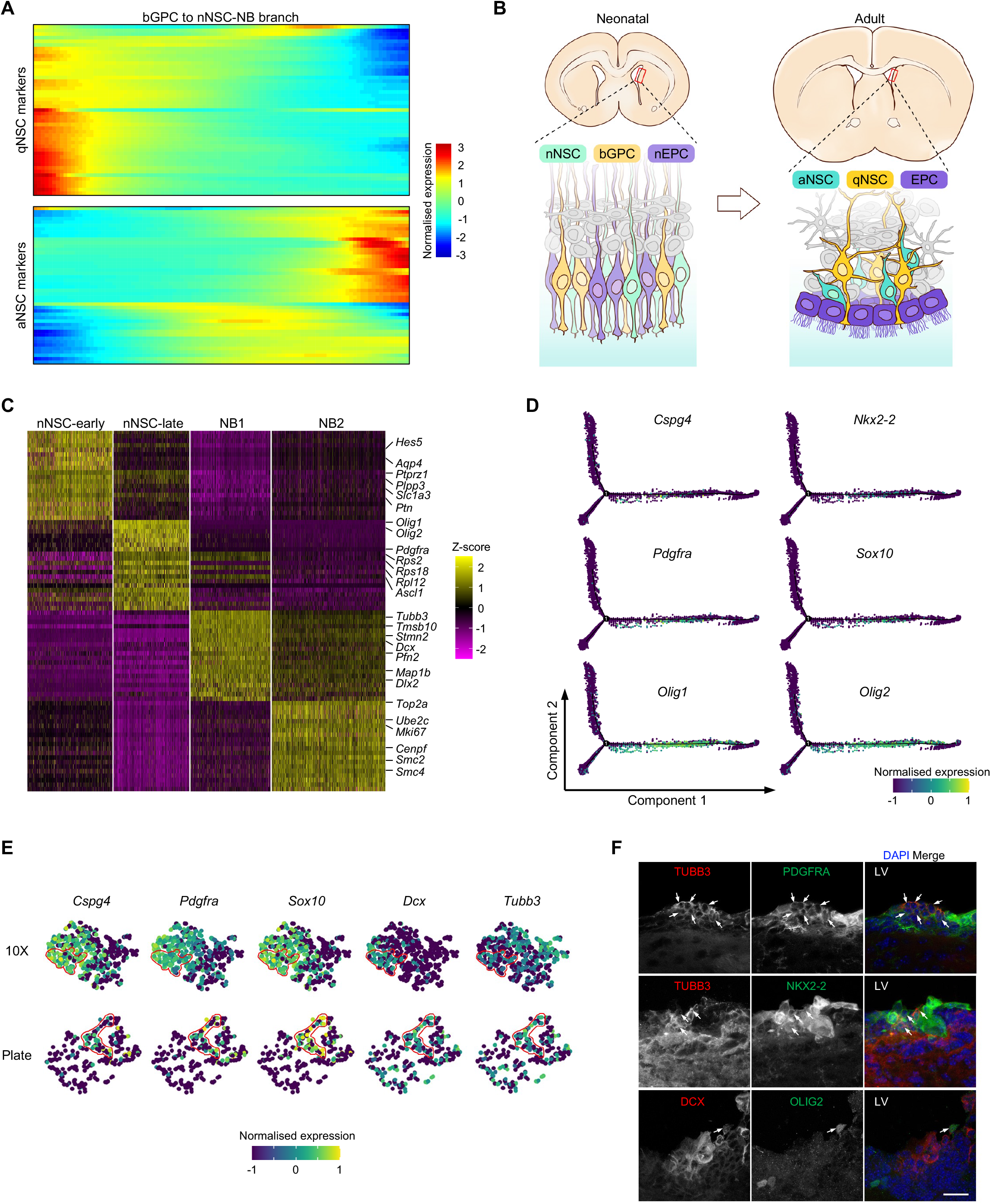
Characteristics of cells along the nNSC-NB branch. **(A)** Heatmaps showing the dynamic expression of marker genes for adult quiescent NSCs (qNSC) and activated NSCs (aNSC) during the differentiation of neonatal bGPCs into nNSCs. **(B)** Schematic diagram illustrating the possible development of the VZ. **(C)** The heatmap of the top 20 signature genes for each of the four clusters along the nNSC-NB branch. Key known marker genes were indicated at the right- hand side. **(D)** Expression profiles of six well-known marker genes of oligodendrocyte progenitor cells (OPCs) along the bifurcating trajectory. **(E)** tSNE visualisations of nNSC_late cells showing co-expressions of OPC markers (*Cspg4*, *Pdgfra*, *Sox10*) and NB markers (*Dcx*, *Tubb3*) from either the 10X or the Plate data. **(F)** OPC-NB bipotent precursors were validated by immunofluorescence of neonatal brain sections. Arrows indicate cells showing positive staining of both the NB marker (TUBB3 or DCX) and the OPC marker (PDGFRA, NKX2-2 or OLIG2). LV, lateral ventricle. The scale bar is 20 μm.

To better characterise the bGPC to nNSC-NB branch, differential expression analysis was performed on the four sequential clusters along the pseudotime branch, including nNSC-early, nNSC-late, NB1 and NB2 (**Figures 1E**). Distinct marker genes were found for each cluster (**Figure 2C**). Gene ontology (GO) enrichment analysis on the differentially expressed genes suggest early-stage nNSCs primarily expressed genes related to gliogenesis (**Supplementary Figure 2A)**. By comparison, late-stage nNSCs exhibited high expression of genes associated with ribosome biogenesis and translation (**Supplementary Figure 2A**), indicating those cells were preparing for extensive protein synthesis. Interestingly, some late-stage nNSCs expressed markers for oligodendrocyte precursor cells (OPCs) (**Figure 2C** and **Supplementary Figure 2A**), which has not been previously reported during neuronal differentiation [33, 34]. This was confirmed by examining the expressions of six OPC marker genes (*Cspg4*, *Nkx2-2*, *Pdgfra*, *Sox10*, *Olig1* and *Olig2*), all of which had high expression levels within cells around the middle of the nNSC-NB branch (**Figures 2D**). Furthermore, some late-stage nNSCs were expressing both OPC markers and NB markers. The observation was seen not only in the 10X data but also in the plate data (**Figure 2E**), thereby ruling out the impact of doublets from droplet-based library preparation. Immunofluorescence analysis of brain sections from neonatal mice confirmed the existence of cells co-expressing NB and OPC markers in the VZ (**Figure 2F** and **Supplementary Figure 2B**). Based on the results, those cells might have dual potentials that can give rise to neurons and oligodendrocytes, and hence we named them OPC-NB bipotent precursors.

The two NB clusters, NB1 and NB2, were located at the end of the nNSC-NB branch (**Figure 1E**). Genes highly expressed in the NB1 cluster were related to neuronal differentiation and the cell cytoskeleton, and NB2 specific genes were mainly involved in cell cycle (**Figure 2C** and **Supplementary Figure 2A**). By combining those results, the sequential molecular events during the differentiation of the nNSC-NB branch can be described as follows: bGPCs first differentiate into early-stage nNSCs that primarily express gliogenic genes; then cells develop into the late-stage nNSCs when genes related to ribosome production and protein translation are upregulated; finally, cells start expressing neuronal markers and cytoskeleton-related genes and are ready to make morphological changes. During the last stage, cells enter mitosis and differentiate into NBs. This differentiation process resembles the transition from qNSCs to aNSCs during adulthood [8–11].

### Multiciliogenesis program dominates bGPC transition into nEPC

Next, the nEPC branch from the bifurcating differentiation trajectory was investigated, consisting of early-stage and late-stage EPCs. Although a set of genes were expressed specifically in the early-stage EPCs, the classic EPC signature genes were only highly expressed in late-stage nEPCs **(Figure 3A)**. GO analysis suggested that genes with higher expression in the early EPCs were mainly associated with glial differentiation and epithelial cell proliferation (**Supplementary Figure 3A**). Cilia-related genes were low during early EPCs but became highly expressed in late EPCs (**Supplementary Figure 3A**).

**Figure 3.**
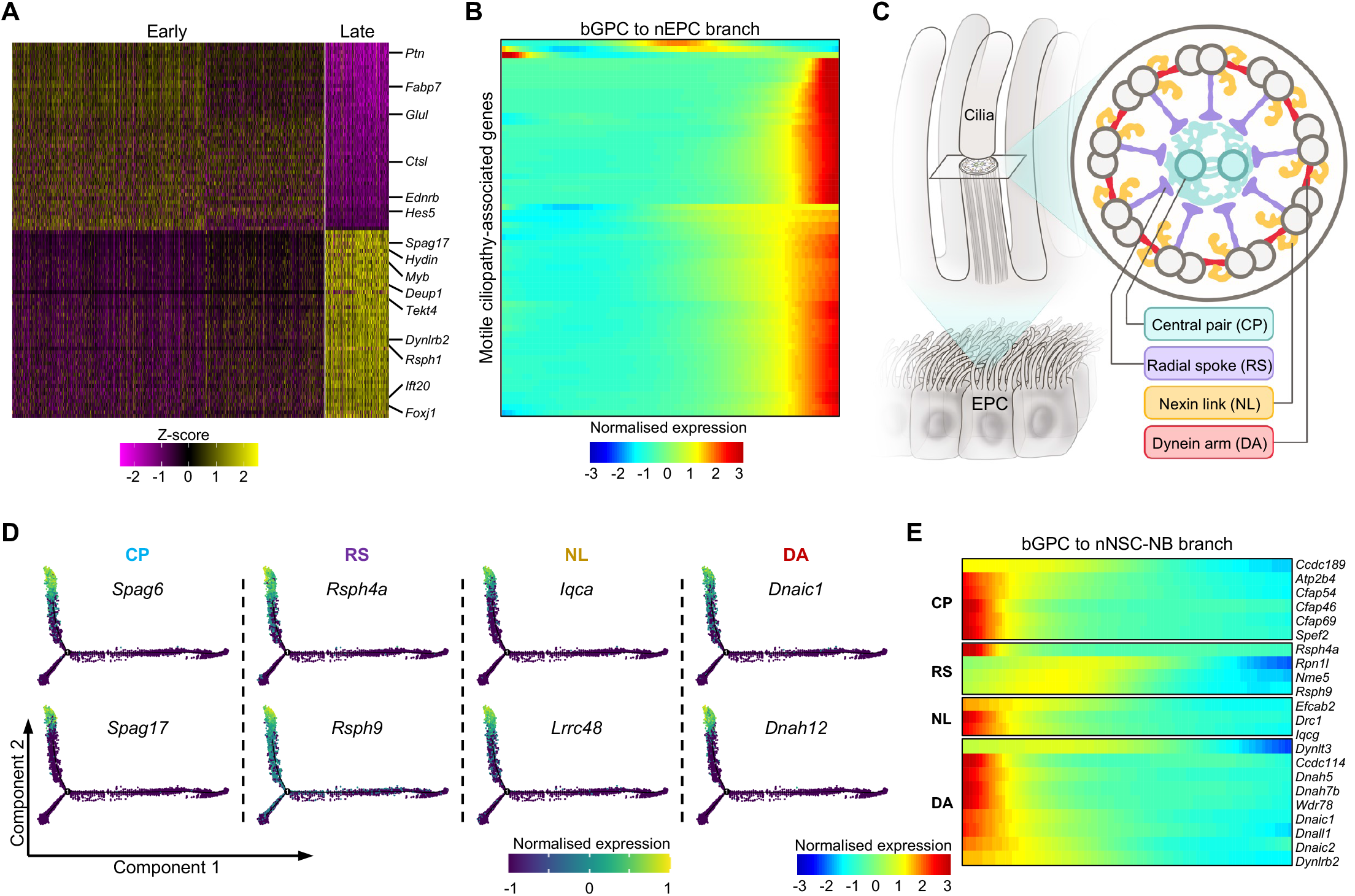
Features of cells along the nEPC branch. **(A)** Heatmap of top 50 marker genes for each of the two EPC clusters. Some key marker genes were indicated at the right-hand side. **(B)** The heatmap showing the dynamic expression of motile ciliopathy-associated genes along the pseudotime trajectory of bGPC to nEPCs. **(C)** Schematic view of the EPC multicilia cross-section. The following key structures are indicated: central pair (CP), radial spoke (RS), nexin link (NL) and dynein arm (DA), all of which are present in multiple motile cilia but not primary cilium. **(D)** The expression profiles of indicated motile-cilia-specific genes along the bifurcating trajectories. **(E)** Heatmap showing the expressions of certain motile-cilia-specific genes are high in bGPCs but become downregulated during the differentiation of bGPCs into nNSC-NBs.

Defects of motile cilia lead to numerous disorders in humans, including hydrocephalus, situs inversus, infertility and respiratory disorders which are collectively termed motile ciliopathies [35, 36]. Currently, 63 genes have been identified to be associated with motile ciliopathies [36]. Among them, 61 were expressed in our dataset and almost all of those genes showed prominent up- regulation along the EPC differentiation branch (**Figure 3B**).

In comparison to primary cilia, motile cilia possess unique structures composed of the central pair (CP), radial spoke (RS), nexin link (NL) and dynein arm (DA) (**Figure 3C)** [35, 37]. Among the generally recognised 26 CP genes [38], 25 were detected in our data; 15 out of 17 RS genes, all 11 NL genes and 39 out of 40 DA genes [39] were detected in our data (**Supplementary Figure 3B**). Almost all of these genes exhibited elevated expression levels during the bGPC- nEPC differentiation (**Figure 3D** and **Supplementary Figure 3B**). The four missing motile cilia structural protein genes were not detected in previously published single-cell transcriptomic datasets from the adult VZ [40] and mass spectrometry data of purified EPC cilia [41] (**Supplementary Figure 3C**), confirming their absence in EPC cilia. These results underscore the robustness of our data.

Of note, we observed extensive expression of many structural proteins of motile cilia in bGPCs. As bGPCs differentiated into nNSCs, the expression levels of these proteins gradually declined (**Figure 3E**). This challenges the notion that structural proteins of motile cilia are exclusively expressed in multiciliated cells, further suggesting a potential role of the down-regulation of ciliary genes during the bGPC-nNSC differentiation.

### Differential analysis on the branches unveils potential hydrocephalus genes impacting EPC differentiation

We further analysed the differentially expressed genes between the bGPC- nEPC and bGPC-nNSC-NB branches to elucidate how the fate of a bGPC was determined. Among the top 3,000 differentially expressed genes, 1,320 genes were specifically up-regulated in the bGPC-nEPC branch (**Supplementary Figure 4A**), and they were termed as nEPC-fate specific genes. GO analysis suggested they were highly enriched in genes related to cilia (**Supplementary Figure 4B**), and 320 of them were classified as ciliary genes (**Supplementary Figure 4C**). The rest of 1000 genes were mainly associated with processes involved in mitochondrial organisation or ATP synthesis (**Supplementary Figure 4D**), which is likely attributed to the enormous energy demand for cilia motility of EPCs [42, 43].

Hydrocephalus is characterised by the enlargement of the brain ventricles caused by obstruction in CSF [19]. To date, more than 400 gene mutations have been identified participating in hydrocephalus formation [19, 44]. Nevertheless, the mechanisms by which most gene mutations lead to hydrocephalus remain elusive. Due to the fact that hydrocephalus is always accompanied by pathological changes in other brain structures, it is challenging to assess the specific impact of a single causal factor. The beating of EPC cilia is the main propelling force for CSF circulation, but only mutations in seven genes, including *Foxj1*, *Mcidas* and *Ccno*, have been identified as causing hydrocephalus by impairing EPC-lineage differentiation [45]. Six of these genes were specifically up-regulated in the nEPC branch (**Figure 4A** and **Supplementary Figure 4E**). Moreover, 76 hydrocephalus-associated gene mutations, previously unreported in the context of ciliogenesis, displayed specific high expression in the nEPC branch (**Figure 4B**). This implies that hydrocephalus in patients carrying these gene mutations is attributed to abnormal EPC-branch differentiation in the VZ.

**Figure 4.**
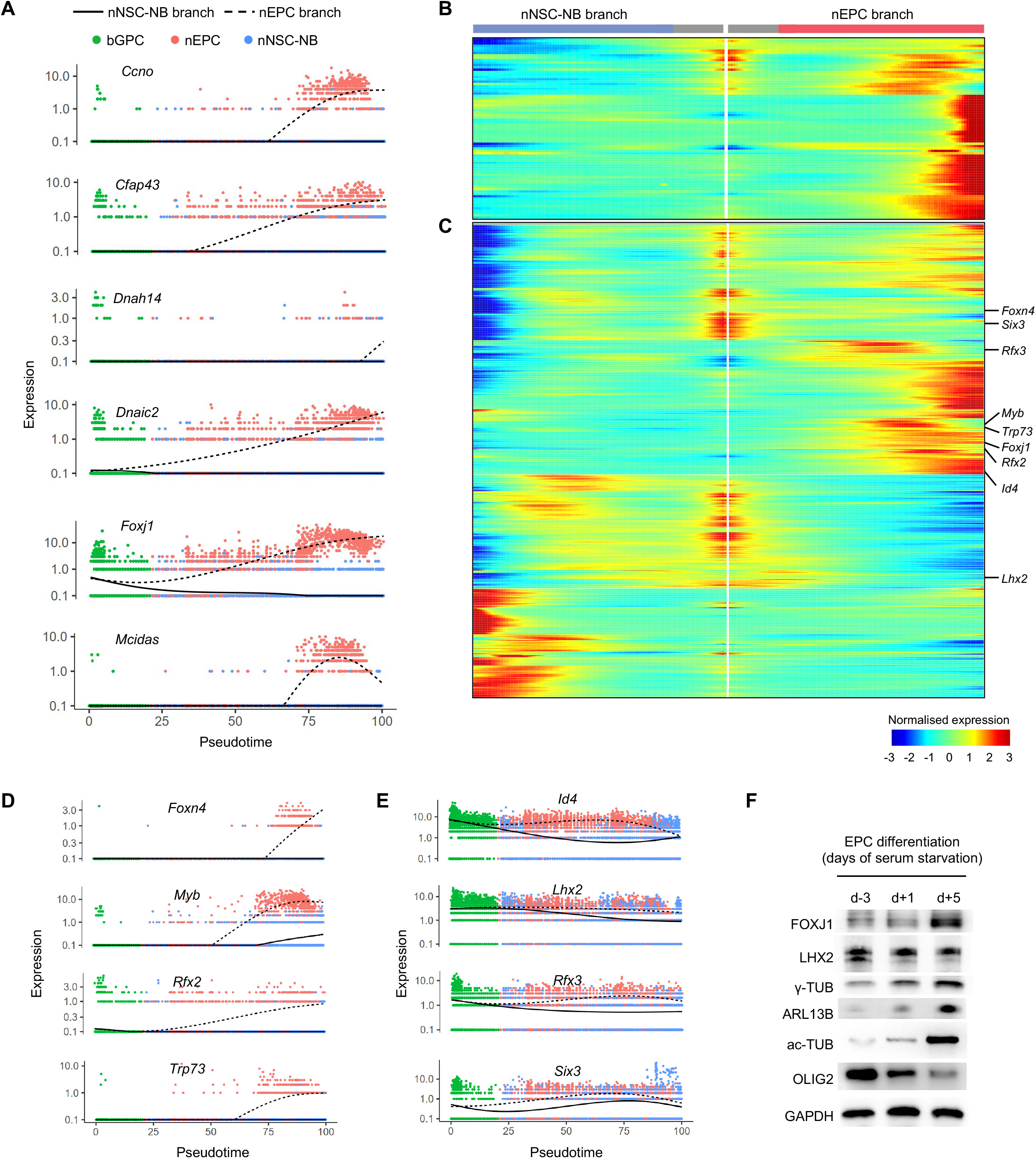
Expression kinetics of hydrocephalus-related genes and EPC-fate regulators in the bifurcating trajectory. **(A)** The six causative hydrocephalus genes were up-regulated specifically in the nEPC branch. **(B)** Heatmap demonstrating 76 genes that have mutations in hydrocephalus were specifically upregulated in the nEPC branch. **(C)** Heatmap showing the dynamic expression of branch-specific differentially expressed transcription factors (TFs). The nine well-known TFs that regulate the EPC-lineage differentiation are labelled at the right-hand side. **(D)** The four known EPC-fate regulators showed elevated expressions specifically in the nEPC branch. **(E)** Expressions of the four known EPC-fate regulators exhibited more pronounced downregulation in the nNSC-NB branch than the nEPC branch. **(F)** Immunoblotting validation of the expression patterns of FOXJ1, LHX2, and OLIG2 during the *in vitro* EPC differentiation. Upregulation of the centriole marker γ-tubulin (γ-TUB), the ciliary membrane marker (ARL13B) and the ciliary microtubule marker acetylated-tubulin (ac-TUB) indicate GPC differentiation into EPCs. GAPDH served as a loading control.

### Transcription factors important for the nEPC fate commitment have distinct expression kinetics in the bifurcating trajectory

Since TF expressions are tightly linked to cell fate determination, we next focused on TFs exhibiting differential expression patterns between the two branches. Among the 311 TFs specifically expressed or up-regulated in nEPCs (**Figure 4C**), *Foxj1* [46–49], *Foxn4* [50], *Rfx2* [48, 51], *Rfx3* [49, 51], *Myb* [52], *Trp73* [53], *Six3* [54], *Lhx2* [55, 56] and *Id4* [57] have been reported to play roles in EPC production or multiciliogenesis. Here, our data uncovered these known EPC-fate regulators could be classified into two categories based on their expression profiles along the bifurcating trajectory. The first category included *Foxj1*, *Foxn4*, *Myb*, *Rfx2* and *Trp73*, which showed progressively increasing expression in the nEPC branch with either no expression or decreased expression in the nNSC-NB branch (**Figures 4D** and **Supplementary Figure 4E**). The second category included *Id4*, *Lhx2*, *Rfx3* and *Six3*, which exhibited relatively even or slightly declined expression pattern in the nEPC branch with a more pronounced down-regulation in the nNSC-NB branch (**Figures 4E** and **Supplementary Figure 4E**).

To experimentally verify the expression patterns of the TFs, RGCs were isolated from the neonatal VZ and differentiated into EPCs *in vitro* under a serum starvation condition [41, 58]. As progenitor cells give rise to EPCs, the expression levels of the centriole marker γ-tubulin (TUBG1 and TUBG2), the ciliary membrane marker (ARL13B) and the ciliary microtubule marker acetylated-tubulin (ac-TUB) increased, while the expression of the OPC marker OLIG2 declined (**Figure 4F**). Thus, the *in vitro* EPC culture and differentiation system faithfully replicated the *in vivo* EPC development process. In agreement with our scRNA-seq data, FOXJ1 expression increased as EPC differentiation progressed, while LHX2 showed no evident changes over time (**Figure 4F**). These results suggested that EPC-fate regulators fulfil mediating EPC branch output through the potential mechanisms of up-regulating their expression in the EPC branch or down-regulating their expression in the NSC branch.

### *Npas1* and *Foxa2* are indispensable for nEPC-lineage differentiation

To identify novel TFs required for the nEPC-branch differentiation from our scRNA-seq data, candidate TFs were depleted by RNA interference (RNAi) in the *in vitro* EPC culture and differentiation system (**Figure 4F**). Multiciliated EPCs were shown to be featured with multiple centrioles or cilia, while other cells contained only two centrioles or one cilium [13, 32, 52]. Therefore, the status of the EPC differentiation could be assessed by immunofluorescent staining of centriole (γ- tubulin) or cilium (ARL13B).

To test whether the *in vitro* system was applicable for the functional assessment of potential TFs, *Mcidas* and *Lhx2*, two genes with explicit effects on EPCs, were selected as positive controls. Consistent with the previous published works [32, 59], *Mcidas* expression peaked in the middle stage of the nEPC lineage (**Supplementary Figure 4E**), and the depletion of *Mcidas* drastically suppressed the conversion of GPCs into EPCs (**Figure 5A** and **B**). In addition, a clear dose-dependent suppression effect of *Mcidas* deficiency was observed. Cells treated with *Mcidas-si2*, which resulted in lower *Mcidas* expression, showed a greater inhibition of multiciliogenesis compared to cells treated with *Mcidas-si1* (**Figure 5A** and **B**). In contrast to *Mcidas*, the loss of *Lhx2* significantly promoted EPC-lineage specification (**Figure 5A** and **B**), which agreed with the *in vivo* results reported earlier [55, 56].

**Figure 5.**
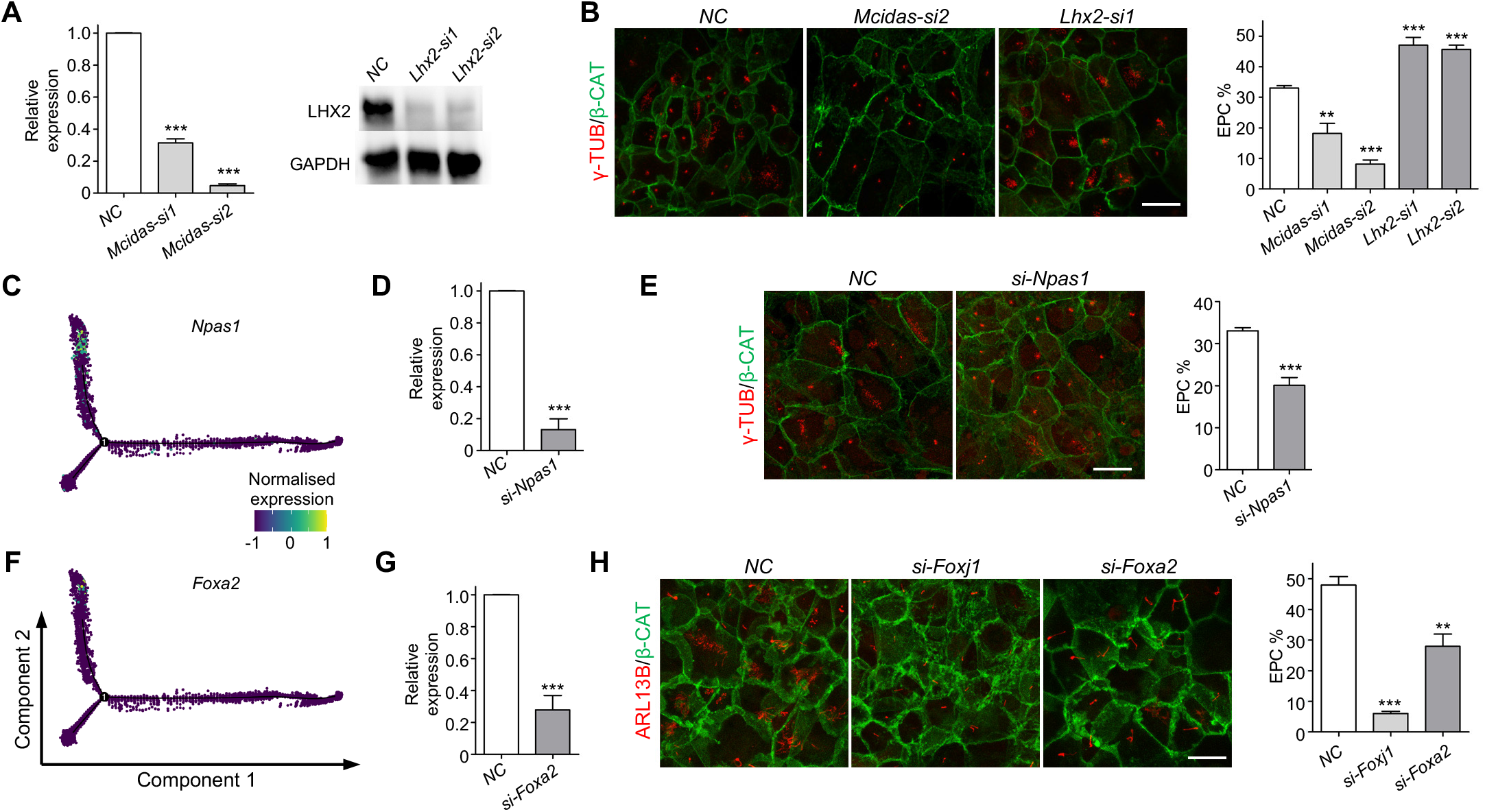
*Npas1* and *Foxa2* facilitate the EPC-lineage differentiation. **(A)** Quantitative PCR (qPCR) and immunoblotting showed efficient depletion of *Mcidas* and *Lhx2* in EPCs by RNAi. NC, negative control. **(B)** Knockdown of *Mcidas* and *Lhx2* promoted and inhibited the EPC production, respectively. Centrioles were labelled by γ-TUB, and cell borders were labelled β-catenin (β- CAT). **(C** and **F)** Expressions of *Npas1* and *Foxa2* were specifically upregulated in the nEPC branch. **(D** and **G)** qPCR showed efficient depletions of *Npas1* and *Foxa2* in EPCs. **(E)** *Npas1* depletion impaired the centriole amplification process during EPC differentiation. **(H)** Knockdown of *Foxa2* compromised the cilium formation process during EPC differentiation. The cilium was labelled by ARL13B. FOXJ1, a well-recognised TF essential for cilium formation, served as positive control. Error bars represent the standard deviation (SD). Asterisks indicate P-values from Student’s t-tests, ** P < 0.01, *** P < 0.001. The scale bars are 20 μm.

Subsequently, we conducted siRNA transfection to deplete several candidate TFs. We first focused on the *Npas1* gene, also known as neuronal PAS domain protein 1, which was found to suppress the differentiation of cortical neurons [60]. Despite its known function, *Npas1* exhibited specific up-regulation in the nEPC branch within the bifurcating trajectory of the VZ development, reaching peak expression in mid-stage nEPCs (**Figure 5C** and **Supplementary Figure 5A**). Moreover, *Npas1* expression in nEPCs was much higher than in glioblasts, NBs and neurons (**Supplementary Figure 5B**), as indicated in the mouse brain development atlas [61], suggesting its involvement in the EPC-lineage differentiation. Deprivation of *Npas1* in the progenitor cells resulted in an evident decrease in the ratio of EPCs from 33.0% to 20.1% (**Figures 5D** and **E**), suggesting that *Npas1* promotes EPC differentiation.

We next investigated the role of *Foxa2*, a member of the Forkhead-box TF family. Similar to the expression patterns of the other two Forkhead TFs *Foxj1* and *Foxn4* (**Figure 4A** and **D**), *Foxa2* also exhibited specific up-regulation in the nEPC branch (**Figure 5F** and **Supplementary** Figure 5C and D). While *Foxa2* depletion did not affect centriole amplification in the early stages of EPC-lineage differentiation (**Supplementary Figure 5E**), it significantly hampered the growth of cilium axoneme in the later stages (**Figure 5G** and **H**), resembling the defects caused by *Foxj1* loss (**Figure 5H** and **Supplementary Figure 5F**) [46, 47]. These results demonstrated that our data successfully identified genes playing critical roles in bGPC fate specification.

### *Tfeb* restrains the EPC-lineage differentiation

Previous studies on the adult VZ have shown that transcription factor EB (TFEB), which modulates lysosome biogenesis, was more abundant in qNSCs compared with aNSCs and facilitated the conversion of qNSCs into aNSCs [62]. In our data, the expression of *Tfeb* in bGPCs was higher than that in nNSCs (**Figure 6A** and **Supplementary** Figure 6A and B). Immunoblotting with an anti-TFEB antibody confirmed that the expression of *Tfeb* was progressively elevated as GPCs differentiated towards the EPCs in the *in vitro* culture system (**Figure 6B**), indicating potential roles for *Tfeb* in EPCs.

**Figure 6.**
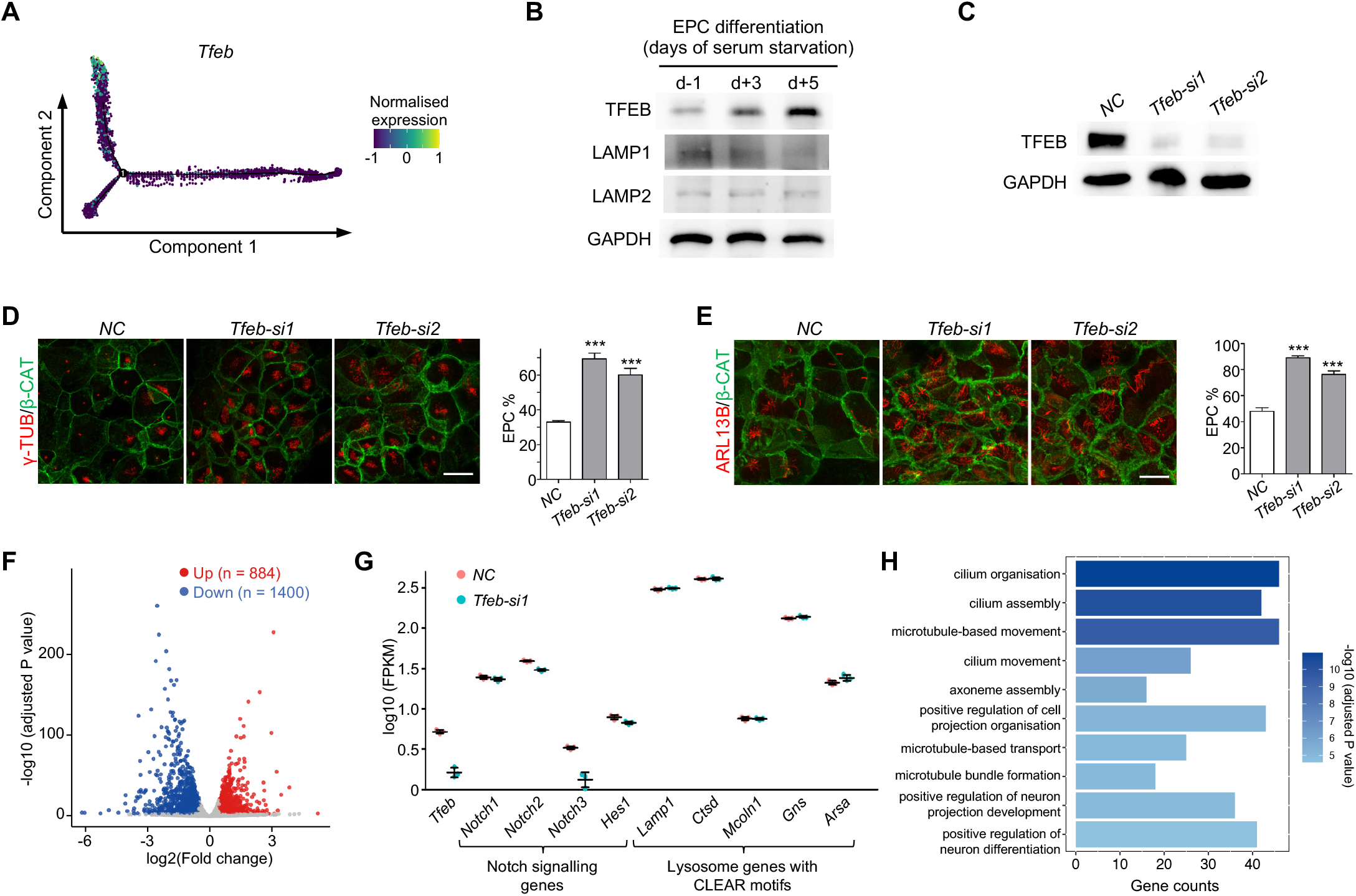
*Tfeb* restrains the differentiation towards EPCs. **(A)** The expression pattern of *Tfeb* along the bifurcating trajectory. **(B)** Immunoblotting of TFEB during the *in vitro* differentiation of GPCs to EPCs. The lysosomal gene LAMP1 and LAMP2 were also shown. GAPDH was used as a loading control. **(C)** Immunoblotting showing efficient depletion of TFEB in EPCs. **(D** and **E)** Immunofluorescence of γ- TUB, indicating centrioles **(D)**, and ARL13B, indicating cilium **(E)**, upon *Tfeb* depletion showing significantly enhanced EPC-lineage differentiation. All of the quantification results above were from three independent experiments. Error bars represent SD. Asterisks indicate P-values from Student’ s t- tests, *** P < 0.001. The scale bars are 20 μm. **(F)** Volcano plot of bulk RNA-seq results showing the significant gene expression changes upon *Tfeb* depletion in EPCs. Red and blue dots indicate significantly upregulated and downregulated genes with adjusted P-value < 0.001, respectively. **(G)** Expression levels of indicated Notch signalling genes and lysosome genes with CLEAR motifs from bulk RNA-seq results. **(H)** Top 10 gene ontology (GO) terms of biological processes from upregulated genes upon *Tfeb* depletion.

Despite *Tfeb* expression was up-regulated as the nEPC-lineage differentiation progressed, the expression of the lysosomal gene *Lamp1* showed a slight reduction and *Lamp2* stayed relatively unchanged (**Figure 6B**). Our scRNA-seq data revealed that the expression of 36 out of the 61 lysosomal genes containing the CLEAR motif [23] declined as bGPCs gave rise to nEPCs (**Supplementary Figure 6C**). These results imply that lysosomal gene expressions in EPCs, somewhat similar to the situation in embryonic stem cells [63], is independent of *Tfeb*.

To further investigate the role of *Tfeb* during EPC differentiation, two independent siRNAs were designed to deplete *Tfeb* in the *in vitro* culture system, both of which achieved efficient depletion of *Tfeb* (**Figure 6C**). Surprisingly, the depletion of *Tfeb* dramatically enhanced EPC differentiation. The examinations on both centriole amplification and cilium formation revealed that the proportion of EPCs increased by nearly two folds (**Figure 6D** and **E**).

Next we performed bulk RNA-seq analysis on *in vitro* differentiated EPCs treated with negative control (NC) or *si-Tfeb*. Three independent biological replicates per group were performed (**Supplementary Figure 6D**). Overall, there were 884 genes that were significantly up-regulated upon *Tfeb* knockdown and 1,400 genes down-regulated (**Figure 6F**). Given that the Notch signalling pathway inhibits multiciliogenesis [64], we examined the expression of genes related to this pathway. In line with data in mammary epithelial cells [65], *Tfeb* loss decreased the expressions of *Notch2*, *Notch3* and *Hes1* (**Figure 6G**), all of which are key components of the Notch signalling pathway. In contrast, *Tfeb* deficiency in EPCs did not affect genes related to lysosome formation (**Figure 6G**). GO analysis of the up-regulated genes revealed that the top five enriched biological processes were all related to cilia (**Figure 6H**), which agreed with the immunofluorescence results (**Figure 6D** and **E**). These findings indicate that *Tfeb* deficiency reduces the expression of the Notch signalling pathway and promotes GPC differentiation towards the EPC. Therefore, it is likely that *Tfeb* plays an inhibitory role in EPC production.

### TFEB activation blocks GPC specification into EPCs through suppressing multicilia-related genes

We next utilised other independent approaches to investigate how *Tfeb* functions in the process of EPC-linage specification. Previous studies have demonstrated that high concentrations of trehalose or sucrose induce lysosome stress, leading to the nuclear translocation and hence the activation of TFEB [23, 66]. Corroborative with the results from the siRNA experiments, 100 mM trehalose or sucrose induced the nuclear localisation of TFEB in GPCs (**Figure 7A** and **Supplementary Figure 7A**), almost abolishing their differentiation into EPCs (**Figure 7B** and **Supplementary Figure 7B**). When GPCs were treated with varying concentrations of trehalose, we observed a strong negative relation between TFEB nuclear translocation and the differentiation efficiency of GPCs into EPCs (**Figure 7A** and **B**). At a concentration of 10 mM trehalose, the nuclear translocation of TFEB could already be observed in many cells (**Figure 7A**). Interestingly, the nuclear localisation of TFEB and the expression of FOXJ1 were mutually exclusive, where TFEB-positive nuclei exhibited no FOXJ1 protein and TFEB-negative nucleus showed positive FOXJ1 staining (**Figure 7C**). The results confirmed the inhibitory effect of TFEB activation on EPC-lineage differentiation. In addition, we found that amino acid starvation also impeded EPC-branch differentiation by inducing TFEB nuclear localisation (**Supplementary** Figure 7C and D).

**Figure 7.**
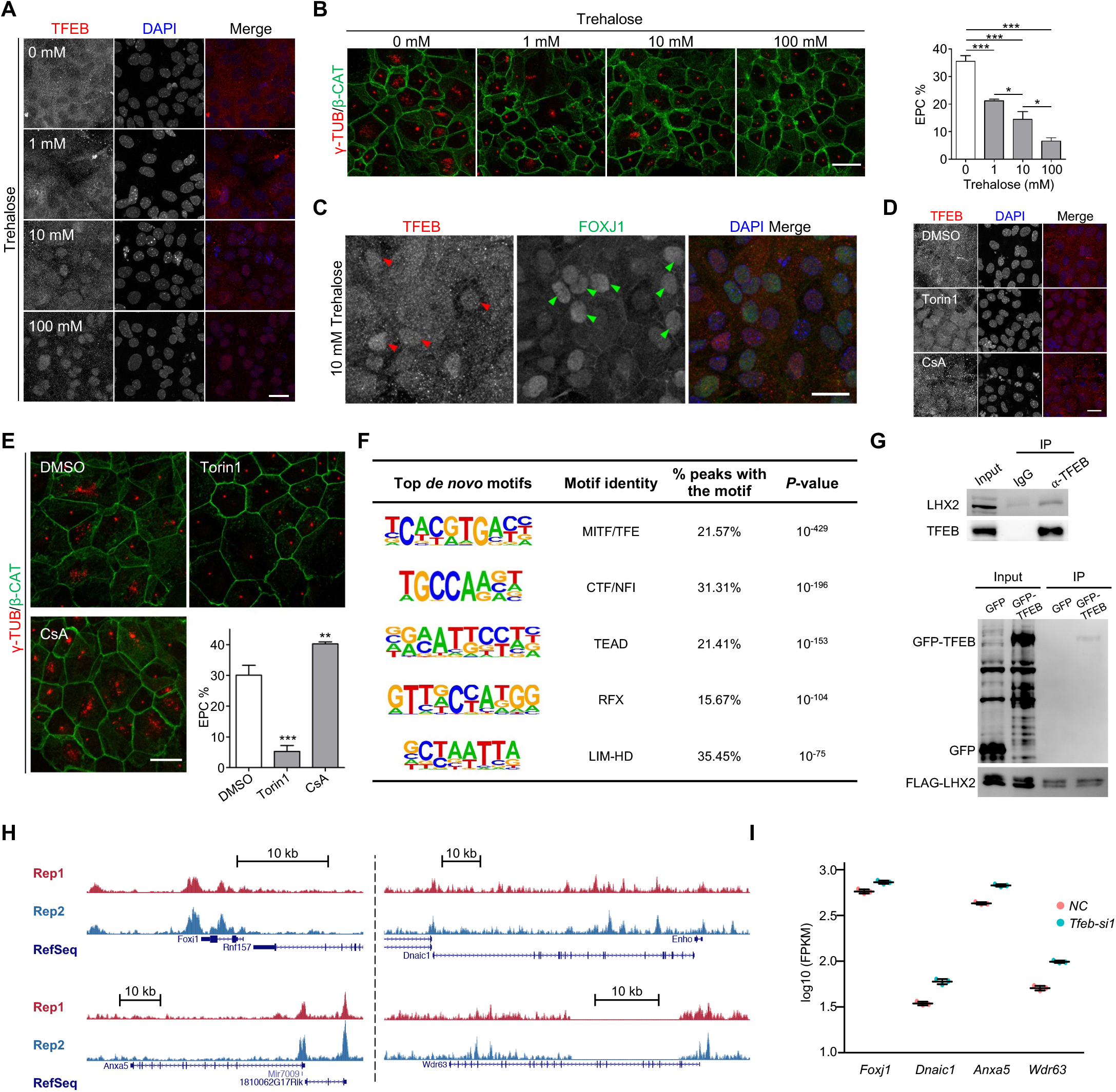
Activated of TFEB cooperates with LHX2 to inhibit multiciliogenesis. **(A)** Nuclear localisation of TFEB by immunofluorescence in the EPC culture system showed a dosage-dependence on trehalose treatment. **(B)** Immunofluorescence of γ-TUB showing the EPC percentage was negatively correlated with the concentrations of trehalose in the culture media. **(C)** Upon 10 mM trehalose treatment, immunofluorescence analyses showing TFEB and FOXJ1 displayed mutually exclusive nuclear localisation in glial cells. Red and green arrowheads denote nuclear localisations of TFEB and FOXJ1, respectively. **(D)** Immunofluorescence analyses showing the translocation of TFEB from the cytosol to the nucleus upon 250 nM Torin1 (mTOR inhibitor) treatment. TFEB remained in the cytoplasm upon 4 μM CsA (Calcineurin inhibitor) treatment. CsA, cyclosporin A. **(E)** Immunofluorescence of γ-TUB showing treatment with Torin1 and CsA impeded and facilitated EPC-lineage differentiation respectively. All of the quantification results from **(B)** and **(E)** were from three independent experiments. Error bars represent SD. Asterisks indicate P-values from Student’s t-tests, *P < 0.05, **P < 0.01, *** P < 0.001. The scale bars are 20 μm. **(F)** Top 5 enriched motifs within TFEB-bound regions identified by ChIP-seq. **(G)** Co-immunoprecipitation revealed the interaction between TFEB and LHX2 in EPCs (top panel) and HEK293T cells (bottom panel). IgG and GFP were used as negative controls. **(H)** UCSC genome browser tracks showing the TFEB bing peaks around four representative ciliary gene loci *Foxj1*, *Dnaic1*, *Anxa5 and Wdr63*. **(I)** Expression levels of four indicated ciliary genes from bulk RNA-seq results.

Prior studies have identified that mTOR phosphorylates TFEB to prevent its activation [67, 68], while calcineurin dephosphorylates TFEB to facilitate its activation [69]. Treatment of GPCs with Torin1, an mTOR inhibitor [67, 68], enhanced TFEB nuclear translocation and hampered EPC differentiation (**Figure 7D** and **E**). On the other hand, treatment with cyclosporin A (CsA), a calcineurin inhibitor [69], slightly increased the efficiency of EPC differentiation (**Figure 7E**), though CsA was unable to induce the nuclear translocation of TFEB (**Figure 7D**). These results implicate that the inhibitory effect of TFEB during EPC-lineage differentiation is closely associated with its phosphorylation.

To further elucidate the molecular mechanisms underlying the role of TFEB in EPC-lineage specification, chromatin immunoprecipitation followed by sequencing (ChIP-seq) was conducted in *in vitro* differentiated EPCs. We identified a total of 10,646 TFEB binding sites in the genome (**Supplementary Figure 7E**). *De novo* Motif analysis returned the top motif that resembled the TFE family motif (**Figure 7F**), confirming the high quality of the data. Interestingly, the RFX family motif and the LIM-HD family motif were also found enriched within the TFEB binding sites (**Figure 7F**). The RFX transcription factors were previously shown to be essential for ciliogenesis [48, 49, 51], and LHX2, a member in the LIM-HD family, were implicated in multiciliogenesis suppression (**Figure 5B**) [55, 56]. It is likely that TFEB cooperates with LHX2 to block the expression of multicilia-related genes. To test this hypothesis, immunoprecipitation showed that both endogenous TFEB in EPCs and exogenous TFEB overexpressed in HEK293T cells interacted with LHX2 (**Figure 7G**). Moreover, TFEB bound to the gene body of *Foxj1* according to our ChIP-seq data (**Figure 7H**), and knockdown of *Tfeb* resulted in elevated expression of *Foxj1* (**Figure 7I**). These results indicated that the binding of TFEB on *Foxj1* might have an inhibitory effect, which explains the immunofluorescence result showing that nuclear translocation of TFEB and FOXJ1 expression were mutually exclusive (**Figure 7C**). Furthermore, TFEB was found to bind to the gene bodies of other cilia-related genes, including *Dnaic1*, *Anxa5* and *Wdr63*, and the binding also seemed to have suppressive effects on their expressions (**Figure 7H** and **I**).

### Overexpression of *Tfeb* induces NSC-lineage differentiation

To exclude the off-target effects of RNAi, an siRNA refractory version of *GFP-Tfeb* was introduced to *si-Tfeb* treated EPCs. The expression of GFP-TFEB significantly mitigated the EPC overproduction caused by the loss of TFEB, as compared to cells expressing GFP only (**Figure 8A** and **Supplementary Figure 8A**). Of note, some GFP-TFEB-positive cells manifested neuron-like morphologies (**Figure 8A**). When GFP-TFEB was overexpressed in wild-type GPCs, 35.4% of them displayed neuron-like morphologies, and only 19.2% gave rise to EPCs (**Figure 8B** and **Supplementary Figure 8B**). In comparison, nearly half of the cells transfected with only GFP differentiated into EPCs, and neuron-like morphologies were rarely observed (**Figure 8B** and **Supplementary Figure 8B**). These findings suggest that an excess of TFEB promotes the differentiation of bGPCs towards the nNSC-NB fate. To probe whether the effects of TFEB overexpression are associated with its transcriptional activity, several constitutively active mutants of TFEB [63] were introduced to GPCs. The mutant GFP-TFEB (S210A) showed the most pronounced nuclear localisation (**Supplementary Figure 8D**). The Ser210 in mouse TFEB was shown to be a conserved residue previously identified as an mTORC1 target [67, 68]. Cells expressing GFP-TFEB (S210A) exhibited a remarkably higher proportion (67.2%) of neuron-like morphologies, and only 3.1% gave rise to EPCs (**Figure 8B** and **Supplementary Figure 8B**). In contrast, cells expressing GFP-TFEB-ΔNLS, a mutant of TFEB in which the nuclear localisation signal (NLS) (amino acids 244 - 247) was mutated to alanines [67], showed a much lower proportion (5.9%) of neuron-like morphologies and a higher differentiation efficiency (35.3%) into EPCs (**Figure 8B** and **Supplementary Figure 8B**).

**Figure 8.**
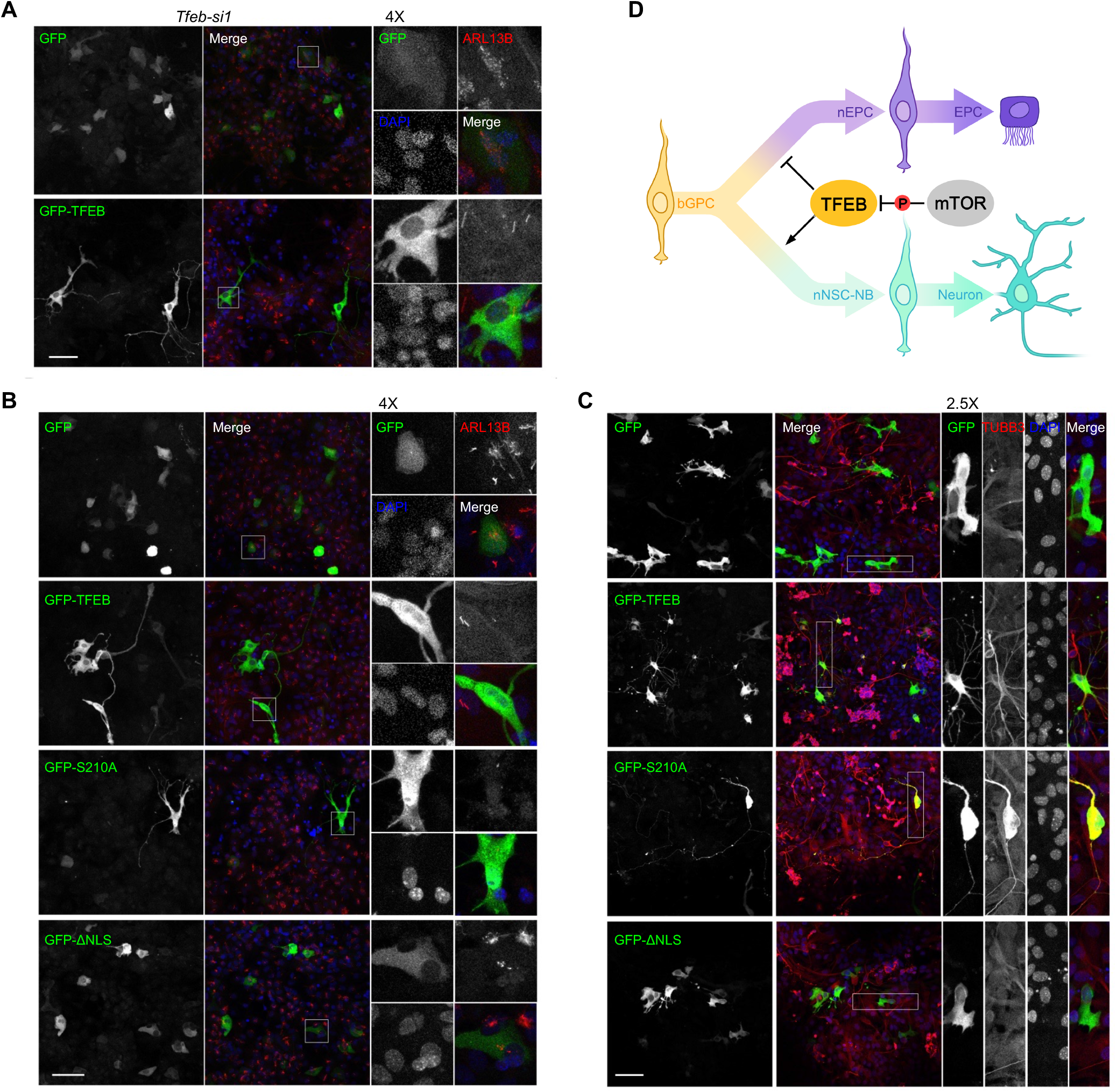
Overexpression of TFEB induces a NSC fate. **(A)** Immunofluorescence of ARL13B showing the overexpression of an RNAi-resistant GFP-TFEB in *Tfeb- si1*- treated GPCs rescued overproduction of EPCs and induced the generation of neuron- like cells. GFP was used as a negative control. The framed region was further magnified to show details. **(B)** Immunofluorescence of ARL13B showing the overexpression of an RNAi-resistant GFP-TFEB and TFEB-S210A, but not TFEB-ΔNLS induced production of neuron-like cells and blocked EPC-lineage differentiation. Cells were serum starved for 5 days and then subjected to immunofluorescence. **(C)** Expression of GFP-TFEB and TFEB-S210A, but not TFEB-ΔNLS induced the conversion of GPCs into neurons. Cells were cultured in the presence of serum. Neurons were labelled by TUBB3 staining. The scale bars are 50 μm. **(D)** A model illustrating how TFEB functions in the process of bGPC specification during VZ development.

To verify that *Tfeb* overexpression in deed promotes the differentiation towards NSCs, RGCs were isolated and cultured *in vitro* in the presence of serum and immunostained with a TUBB3 antibody. 39.4% of GFP-TFEB-positive cells and 30.9% of GFP-TFEB (S210A)-positive cells were differentiated into neurons (**Figure 8C** and **Supplementary Figure 8C**). In contrast, the ratio fell to only 6.4% and 11.9% in GFP-positive and GFP-TFEB-ΔNLS-positive cells, respectively (**Figure 8C** and **Supplementary Figure 8C**). In addition, 33.7% of GFP-TFEB-positive cells and 48.8% of GFP-TFEB (S210A)-positive cells displayed neuron-like morphologies without expressing TUBB3 (referred to as neuron-like cells), compared to only 11.3% in GFP-positive and 10.3% in GFP- TFEB- ΔNLS- positive cells ( **Figure 8C** and **Supplementary Figure 8C**).

Similar to the situation where TFEB activation in the adult VZ promotes the transition of qNSC to aNSC, the TFEB activation in neonatal VZ facilitates bGPC specification towards the nNSC-NB lineage. Collectively, these findings indicate that TFEB controls neonatal EPC/NSC bifurcation. Activated TFEB, on one hand, prevents the overproduction of EPCs by interacting with LHX2 and jointly binding to the regulatory regions of ciliary genes such as *Foxj1* to suppress their expressions. On the other hand, activated TFEB promotes the differentiation of bGPCs into nNSCs, a process that can be modulated via the phosphorylation on the S210 residue by mTORC1 (**Figure 8D**).

## Discussion

### Unveiling the co-differentiation lineage roadmaps of postnatal EPCs and NSCs

Previous transcriptomics studies on postnatal NSCs have predominantly focused on the transition of qNSCs to aNSCs in the adulthood [8–11], without defining the developmental origins of these two NSC subtypes and their primary niche EPCs. In this study, we conducted scRNA-seq analysis of the VZ in neonatal mouse brains and identified three main states of RGCs, namely bGPC, nNSC-NB and nEPC (**Figure 1B**, **C**, **Supplementary** Figure 1C and D). We described the bifurcating trajectory of bGPC differentiation towards nNSC-NB and nEPC (**Figure 1E**). By integrating our data with the transcriptomic profiles of NSCs at the adult stage, we extrapolated that neonatal bGPCs become qNSCs in adulthood, while nNSCs evolve into aNSCs.

Our data demonstrated that the developmental processes of postnatal EPCs and NSCs are connected. For the NSC-lineage specification, the expression of specific ciliary structural proteins needs to be down-regulated from bGPCs (**Figure 3E**). At the same time, certain TFs important for multiciliogenesis remain relatively unchanged along the EPC differentiation, but their expressions exhibit a decline along the NSC lineage (**Figure 4E**). These findings unveil the diverse gene regulatory mechanisms for the cell fate decision exploited by bipotent progenitors during their differentiation processes.

### Identification of OPC-NB bipotent precursors

It has been previously assumed that OPCs and NBs differentiate along separate lineages [33]. However, by examining the expression profiles of marker genes and performing immunofluorescence staining, we observed that intermediate state cells in the NSC branch exist, where both OPC and NB marker genes are simultaneously expressed in those cells (**Figures 2E** and **F**). This observation suggests the existence of a bipotent progenitor cell state in the VZ capable of participating in both oligodendrocyte and neuronal differentiation. Future studies should focus on verifying the differentiation capacity of these OPC-NB bipotent precursors. Given that oligodendrocytes are responsible for ensheathing axons to ensure the smooth conduction of action potentials [70], our findings point to the possibility of new avenues for regenerative approaches to restore myelin integrity and promote functional recovery in cases of neural injuries. By manipulating the fate determination of these bipotent precursors, it may be possible to enhance the production of both oligodendrocytes and neurons, leading to improved repair outcomes.

### A resource for systematic identification of fate determinants for bGPC differentiation

Through differential analysis of gene expressions in the two branches from the bifurcating trajectory, we identified several TFs that have not been characterised in the context of neonatal gliogeneis may play roles in fate determination towards EPCs or NSC-NBs (**Figure 4C**). Experimental validations revealed that *Npas1* and *Foxa2* enhance the EPC-lineage differentiation by facilitating centriole amplification and ciliogenesis, respectively (**Figure 5E** and **H**). Interestingly, *Tfeb* emerged as a key player in both EPC-lineage and NSC-lineage specifications, governing the fate of bGPCs (**Figure 8D**). These results demonstrate the power of our scRNA-seq data and the potential of this resource for understanding the molecular mechanisms underlying bGPC fate determination. Further experimental validation of other potential TFs may help understanding the fate-determining nodes of bGPCs and provide valuable insights into the development and function of the VZ.

### TFEB activation restrains excessive multiciliogenesis

Strict control over the number and size of organelles is a prerequisite for maintaining cell homeostasis due to the limited availability of cellular components [71, 72]. In particular, multiple motile cilia, a hair-like organelle protruding from the cell surface, requires vast material consumption for their formation and substantial energy expenditure for beating [42, 43]. Therefore, understanding the mechanisms that restrain excessive muliticilia generation is critical. We found that, during the differentiation of bGPCs into multiciliated EPCs, the expression of *Tfeb* was up-regulated both *in vivo* and *in vitro* (**Figure 6A** and **B**). To our surprise, *Tfeb* depletion further facilitated EPC production (**Figure 6D** and **E**), while activating TFEB by means of lysosome stress (**Figure 7A, B, Supplementary** Figure 7A and B), amino acid starvation (**Supplementary** Figure 7C and D), or mTOR inhibition (**Figure 7D** and **E**) markedly blocked EPC generation. There was a strong negative correlation between forced TFEB activation and the differentiation of progenitors towards multiciliated EPCs (**Figure 7A**, **B** and **C**). Currently, the experimental data from our study suggest that the expression level of *Tfeb* is important for the proper differentiation of bGPCs. Under the normal developmental condition, the expression of *Tfeb* goes up in the EPC branch, acting as a “brake” to prevent excessive multiciliogenesis. Without *Tfeb*, bGPCs would predominantly differentiate into EPCs, and hence negatively impact the differentiation into NSC-NBs.

Furthermore, we uncovered that TFEB cooperates with LHX2 (**Figure 7G**), a TF known to suppress multicilia formation, and binds to many multicilia-related genes such as *Foxj1* and *Dnaic1* (**Figure 7H**), thereby suppressing their expressions to prevent excessive multiciliogenesis. Our study elucidates the molecular mechanisms through which TFEB mediates multicilia formation and highlights the importance of TFEB in maintaining proper cellular homeostasis.

### Targeting *Tfeb* to control postnatal NSC production could be a potential strategy for the prevention and treatment of NDDs

The major hallmarks of NDDs are accumulation of disease-associated proteins and extensive loss of neurons [73]. Recent studies have shown that *Tfeb* plays crucial roles in the NDD pathogenesis. Dysregulation of *Tfeb* expression, nuclear localisation and transcriptional activity has been observed in various NDDs, including Alzheimer’s Disease, Parkinson’s disease and Huntington’s disease [74]. The activation of *Tfeb* has been widely proven to ameliorate the pathological protein aggregates in neurons [66, 74–76]. It promotes the removal of these aggregates through the up-regulation of lysosomal biogenesis and autophagy. Notably, the clinical drugs Aspirin and Rapamycin have shown promising results by enhancing TFEB activity and reducing protein aggregation in disease models [76–79].

In this study, we uncovered that *Tfeb* dictates the balance of neonatal EPC/NSC bifurcation in the developing VZ. Activation of *Tfeb* impedes the expression of cilia-related genes (**Figure 7I**), thus suppressing the differentiation of GPCs towards EPCs while promoting their differentiation towards NSC-NBs. Therefore, utilising approved *Tfeb*-targeted agonists to modestly increase NSC production in the developing VZ may potentially compensate for neuronal loss in NDDs, leading to preventive or delaying effects on these conditions. Further investigations are needed to fully understand the molecular mechanisms underlying *Tfeb*-mediated NSC production and to optimise the therapeutic strategies.

## Methods

### Antibodies and oligonucleotide sequences

Antibodies used for immunoblotting or immunofluorescence can be found in **Supplementary Table S1**. Sequences of siRNA and qPCR primers can be found in **Supplementary Tables S2 - 4**.

### Mice

Mice were maintained in laboratory animal center at Southern University of Science and Technology. Procedures were approved by Experimental Animal Welfare Ethics Committee, Southern University of Science and Technology.

### RGC isolation

Dissection of VZs from neonatal mouse brains was performed as described [58]. Briefly, the midbrain, the olfactory bulb, meninges and the hippocampus were removed in sequence to attain the telencephalon. Then VZ-containing tissues were separated from the telencephalon. Tissues from 4 mice were pooled, transferred to a 1.5-mL Eppendorf tube, and digested with freshly prepared Papain solution (10 U/mL, Worthington, cat. no. LS003126) at 37°C for 30 minutes. Then the Papain solution was aspirated, and 1 mL culture medium, which consisted of DMEM (Gibco, cat. no. SH30256.01) supplemented with 20% FBS ( Gibco, cat. no. 30044333 ), 100U/ ml penicillin and 100 μg/mL streptomycin sulfate (Hyclone, cat. no. SV30010), was added and incubated for 1 minute to terminate the digestion. The supernatant was removed, and another 1 mL culture medium (20% FBS) was added. The digested tissues were gently pipetted ups and downs 20 times using a P1000 tip to dissociate the cells, and then cells were spun down in a centrifuge at 350 RCF for 5 minutes. The supernatant was discarded, and 1 mL blocking buffer (1% BSA, Sigma, cat. no. V900933 in PBS, Hyclone, cat. no. SH30256.01) was used to resuspend the cell pellet. Cells were spun down again at 350 RCF for 5 minutes and the supernatant was discarded. 200 μL blocking buffer containing 2 μL CD133- PE antibody (eBioscience, cat. no. 12-1331-82) or the IgG isotype control (eBioscience, cat. no. 12-4301-82) was added to resuspend the cell pellet and label the cells on ice for 30 minutes. Cells were then washed once with 1 mL blocking buffer and finally resuspended in 1 mL blocking buffer containing 1 μg/mL DAPI (Sigma, cat. no. D8417). The cell suspension was transferred to a FACS tube for sorting. CD133-positive, DAPI-negative single cells were collected into a 15 mL tube or sorted into each well of 384-well plates for scRNA-seq library construction.

### Single-cell RNA-seq library construction

For the droplet-based scRNA-seq experiment, RGCs from P0 and P5 were used. The V3 chemistry of the 3’ Single Cell Gene Expression kit was used according to the 10X Genomics user guide CG000204 Rev D. For the plate-based scRNA-seq experiments, two replicates of RGCs from P0 were used. The experiments were performed exactly according to the step-by-step protocol described previously [29], where the cDNA was amplified for 11 cycles, and the final library was amplified for 11 cycles.

### Single-cell RNA-seq data analysis

For scRNA-seq data, reads were processed using the STARsolo [80] pipeline as previously described [29]. FastQ files generated from Illumina NovaSeq 6000 were aligned to the mm10 mouse reference genome, and the gene expression matrix containing the UMI count for each gene in each cell was obtained. This output was imported into the R toolkits for downstream analyses. Cell doublets in 10X data were identified and filtered using the DoubletFinder package [81]. Samples from P0 and P5 stage in the 10X data or samples from different plates in the plate data were merged using Seurat [82]. The top 2000 highly variable genes were obtained and used for the principal component analysis (PCA). The first 20 PCs were used for graph-based clustering to identify distinct groups of cells. These groups were projected onto 2D tSNE plane. The endothelial cells, microglia and pericytes/fibroblasts, which were annotated by the SingleR package [83], were not included in the analysis. Dying cells, which showed low number of detected genes and UMIs and high mitochondrial genes were also excluded from the subsequent analyses. The remaining cells were re-clustered and the variable genes were identified as ordering genes using the ‘differentialGeneTest’ function in the Monocle2 package [30]. Then dimensionality reduction was done through the DDRTree method and the trajectory of cellular differentiation was constructed using the ‘orderCells’ function. Pseudotime values were acquired by setting the bGPC state as the root. Differential expression analyses to identify the cluster markers and the branch markers were performed using the ‘FindAllMarkers’ function in Seurat and the ‘BEAM’ function in Monocle2, respectively. Trajectory analysis using the diffusion map algorithm was completed with the destiny package [31]. GO analysis for enriched biological processes was performed using clusterProfiler package [84]. The exact code and parameters of the procedures can be found in the GitHub repository: https://github.com/sibszheng/VZ_development.

### *In vitro* culture of EPCs

*In vitro* cultured EPCs were obtained as described [58], with some modifications. The VZs from 4 mice were pooled, transferred to a 1.5 mL Eppendorf tube, and digested with freshly prepared Papain solution at 37°C for 30 minutes. Then the 10 U/mL Papain solution was aspirated, and 1 mL culture medium (20% FBS) was added, followed by 1 minute of incubation to terminate the digestion. The supernatant was removed, and another 1ml culture medium (20% FBS) was added. The digested tissues were gently pipetted up and down by 20 times with a P1000 tip to dissociate the cells mechanically. A 25 cm^2^ flask was pre-coated with 5 μg/mL Laminin (ThermoFisher, cat. no. 23017015) in PBS for 8 - 12 hours, rinsed twice with PBS, and then 3 mL culture medium (20% FBS) was added. The cells were seeded into the flask and cultured at 37°C in an atmosphere containing 5% CO_2_. After 24 hours, the old medium was replaced by fresh culture medium (20% FBS). After another 24 hours, the old medium was aspirated, and the flask was vigorously shaken to remove the neuroblasts and neurons. The remaining radial glia-enriched cells were rinsed once with PBS and further cultured to 30 - 40% confluency. Then the culture medium was aspirated, washed once with PBS, and 1 mL of 0.05% Trypsin ( Gibco, cat. no. 25300062) was added for digestion at 37 °C for 5 minutes. 2 mL culture medium (20% FBS) was added to terminate the digestion. The cells were transferred to a 15ml tube and centrifuged at 900 RPM for 5 minutes. The supernatant was discarded, and 1 mL culture medium (20% FBS) was added to resuspend the cell pellet. 29-mm glass-bottom dishes (Cellvis, cat. no. D29-14-1.5-N) were pre-coated with 5 μg/mL Laminin in PBS for 8 - 12 hours and rinsed twice with PBS before use. 250 μL of cell suspension was transferred into each well of the dish and incubated at 37°C until the cells were completely adhered. After adding 1 mL culture medium (10% FBS), the cells were further cultured to 100% confluency, and then maintained in starvation medium (culture medium without FBS) to induce differentiation into EPCs.

### Plasmids

The full-length *Tfeb* ( NM_001161722) and *Lhx2* ( NM_010710) were amplifi ed by PCR from total cDNAs of mouse EPCs and constructed into pLV-EGFP-C1 and pcDNA3.1-FLAG to express GFP-tagged and FLAG-tagged fusion proteins, respectively. The cDNAs for the S210A mutant (TCC → GCC), the ΔNLS mutant (1032 AGAAGACGCAGG → GCAGCAGCCGCG) and the RNAi-resistant constructs of *Tfeb* (956 GCGAGAGCTAACAGATGCT → AAGGGAATTGACTGACGCA) were produced by PCR. All the constructs were verified by sequencing.

### siRNA and plasmid transfection

For RNAi, siRNAs were transfected into cultured RGCs/EPCs using Lipofectamine RNAiMAX (ThermoFisher, cat. no. 13778150) at serum starvation day -1 (SS d-1) and day 2 (SSd+2), respectively. For one 29-mm dish, the original medium was discarded and replaced by 500 μL fresh culture medium (10% FBS) or starvation medium before transfection. 2 μL of 20 μM siRNA (Genepharma) were mixed in 125 μL Opti-MEM (Gibco, cat. no. 31985070) by vortexing. 3 μL Lipofectamine RNAiMAX were mixed in 125 μL Opti-MEM by vortexing and incubated at room temperature for 5 minutes. The two diluents were then mixed by vortexing and incubated at room temperature for 20 minutes. Finally, the mixture complex was added to the culture medium (10% FBS) or starvation medium. After 24 hours of transfection, cells were rinsed with PBS twice and serum-starved for centriole staining at SS d+2 or cilium staining at SS d+5.

For the expression of GFP-tagged proteins, cultured RGCs were transfected using Lipofectamine 2000 (ThermoFisher, cat. no. 11668019) at SS d-1. For one 29- mm dish, the original medium was discarded and replaced by 500 μL fresh culture medium (10% FBS) before transfection. 2 μg plasmid were mixed in 125 μL Opti- MEM by vortexing. 1.5 μL Lipofectamine 2000 were mixed in 125 μL Opti-MEM by vortexing and incubated at room temperature for 5 minutes. Then the two diluents were mixed together by vortexing and incubated at room temperature for 20 minutes. Finally, the mixture complex was added to the culture medium. After 24 hours of transfection, cells were rinsed with PBS twice, subjected to serum starvation to observe EPC-lineage differentiation at SS d+5 or maintained in culture medium (10% FBS) for NSC-lineage differentiation assay.

For the rescue experiments, cultured RGCs were transfected using Lipofectamine 2000 at SSd-1 to express RNAi-insensitive GFP-TFEB and GFP on top of the transfection of siRNAs. For the co-immunoprecipitation experiments, HEK293T cells were transfected using Lipofectamine 2000 to express the exogenous proteins.

### Drug treatment

Cultured RGCs were incubated with 1 - 100 mM trehalose (Selleck, cat. no. S3992), 100 mM sucrose ( Sigma, cat. no. V900116), 250 nM Torin1 ( Selleck, cat. no. S2827), or 4 μM cyclosporin A (Selleck, cat. no. S2286) in starvation medium from SS d0 to SS d+2. For amino acid starvations, cells were cultured in DMEM without amino acids (Wako, cat. no. 048-33575) from SS d0 to SS d+2. Cells were fixed and subjected to immunostaining after drug treatment.

### Immunofluorescent staining of cultured cells

Cells were fixed with 4% paraformaldehyde (PFA) in PBS for 15 minutes at room temperature and permeabilised with 0.5% Triton X-100 for 15 minutes. For centriole staining, cells were pre-extracted with 0.1% Triton X-100 for 30 seconds before fixation. After 1-hour blocking at the room temperature with the blocking buffer (1% BSA in PBS), the samples were labelled with primary antibodies (diluted in the blocking buffer) and incubated overnight at 4°C. Subsequently, the samples were rinsed three times with the blocking buffer, followed by incubation with secondary antibodies and 1 μg/mL DAPI (diluted in the blocking buffer) for 1 hour at the room temperature. The samples were rinsed three times with PBS and then mounted using an anti-fade mounting medium (Dako, cat. no. s3023). Multi-layered confocal images were captured by using Leica TCS SP8 system and processed with maximum intensity projections.

### Tissue sectioning and immunostaining

Cryo-sectioning was employed for 20-μm-thick tissue sections. Prior to the dissection of the brains, neonatal mice were perfused with PBS followed by 4% PFA. The brains were then postfixed in 4% PFA at 4°C for 4 hours. The fixed brains were soaked overnight in 30% sucrose at 4°C for dehydration. Next, the brains were embedded in OCT compound (Leica, cat. no. 14020108926) and coronally cryo- sectioned using a Leica CM1950 cryostat microtome. The sections were collected onto glass slides.

To perform immunostaining, the cryo-sections were first rinsed with PBS to eliminate the OCT compound. Subsequently, the sections were incubated in blocking buffer (10% normal goat serum and 0.3% Triton X-100 in PBS) for 1 hour at room temperature to prevent non-specific binding. Next, the sections were labeled with primary antibodies in the blocking buffer overnight at 4°C. After three rinses with the blocking buffer, the samples were incubated with secondary antibodies and 1 μg/mL DAPI in the blocking buffer for 1 hour at room temperature. The sections were then rinsed three times with PBS and mounted in anti-fade mounting medium.

### ChIP-seq

ChIP-seq experiments were performed using the ChIPmentation protocol as previously described [85]. Briefly, 5 × 10^6^ EPCs at SS d+5 were collected and crosslinked with 1% formaldehyde in PBS. Then the crosslinking was stopped by adding glycine to a final concentration of 125 mM. The cells were washed twice with PBS and resuspended in 300 μL sonication/IP buffer. The chromatin was fragmented by sonication using a Bioruptor Pico for 4 minutes (30 seconds on, 30 seconds off). The sonicated chromatin was centrifuged at 16,000 RCF for 10 minutes at 4°C. The supernatant was incubated with 10 μL protein A Dynabeads pre-bound with 1 μg anti-TFEB antibody on a rotator overnight in a cold room. Then the IP was washed once with the RIPA buffer, once with the low salt buffer, once with the high salt buffer and twice with 10 mM Tris-HCl, pH 8.0. The beads were resuspended in 30 μL tagmentation mix containing 1 μL Tn5 and incubated on a thermomixer at 37°C for 5 minutes. Finally, the beads were washed twice with the low salt buffer and once with TE. Subsequently, the beads were resuspended in 100 μL ChIP elution buffer and incubated at 65°C overnight for the reverse crosslinking. The tagmented DNA was purified using the ZYMODNA Clean & Concentrator-5 kit. The library was prepared by using standard Nextera PCR primers.

### Bulk RNA-seq library construction

At SS d+5, total RNA of EPCs treated with NC or *Tfeb-si1* were extracted using the RNAsimple Total RNA Kit (TIANGEN, cat. no. DP419). Library construction was performed using the SHERRY protocol as described previously [86]. Briefly, the amount of RNA was quantified using a Nanodrop. 2 μg total RNA were mixed with 2 U RNase-free DNase and incubated at 37°C for 30 minutes. The RNA was purified by 2 × VAHTS DNA Clean beads. Then 200 ng purified RNA was used for reverse transcription using 100 U Maxima H Minus Reverse Transcriptase. The resulting RNA/cDNA hybrid was tagmented with 2.5 μL Tn5. The tagmented product was purified using 2 × VAHTS DNA Clean beads. Library preparation was done using standard Nextera primers.

### ChIP-seq and bulk RNA-seq data analysis

For ChIP-seq data, reads were mapped to the mm10 mouse reference genome using hisat2 [87]. Then the reads with mapping quality less than 30 were removed by samtools [88] and deduplicated using the ‘ MarkDuplicates’ function from the Picard tool (https://broadinstitute.github.io/picard/). Peaks were called on the output BAM file using MACS2 [89]. BedGraph files generated from MACS2 callpeak were converted to bigWig files and visualised via UCSC genome browser [90]. Two biological replicates were performed. Only peaks that were present in both biological replicates were considered. *De novo* Motif discovery was performed using findMotifsGenome.pl in the HOMER suite [91] on the narrowPeak files returned from MACS2.

For bulk RNA-seq data, reads were mapped to the mm10 mouse reference genome using hisat2 and supplied with gene annotation from GENCODE vM25 [92]. Gene expression was quantified by HTSeq [93] and differential gene expression analysis was conducted using DESeq2 [94] package.

The exact code and parameters of the procedures can be found in the GitHub repository: https://github.com/sibszheng/VZ_development.

### Immunoblotting

Cultured cells were directly lysed in 1 × SDS-PAGE loading buffer (Biosharp, cat. no. BL502B) and boiled at 99°C for 10 minutes. Proteins separated by SDS-PAGE (Sangon, cat. no. C681102) were transferred to nitrocellulose membranes. Blots were blocked with 3% BSA diluted in TBS ( Sangon, cat. no. B548105) with 0.05% Tween- 20 ( TBST) for 1 hour at room temperature and then incubated with primary antibodies (diluted in 1% BSA in TBST) at 4°C overnight. After extensive rinse with TBST, membranes were incubated with secondary antibodies (diluted in 1% BSA in TBST) at room temperature for 1 hour. After thorough wash in TBST, protein bands were visualised with enhanced chemiluminescent reagent (Bio-Rad, cat. no. 1705061) and exposed to Tanon 5200 Chemiluminescent Imaging System.

### Immunoprecipitation

Cells were lysed in Sonication/IP Buffer containing protease inhibitor cocktail (1:200, Abcam, cat. no. ab201111) and phosphatase inhibitor cocktail (1:100) (Sigma, cat. no. P0044). Without fixation and sonication, the lysate was centrifuge at 16,000 RCF for 10 minutes at 4°C, and the supernatant was incubated with antibody-bound beads overnight at 4°C. After beads were washed once with RIPA Wash Buffer, once with Low Salt Wash Buffer, once with High Salt Wash Buffer, once with LiCl Wash Buffer and twice with 10 mM Tris-HCl (pH 8.0), proteins associated with the beads were eluted in 1 × SDS- PAGE loading buffer at 99 °C for 10 minutes. 5 × 10^6^ cells and 1 μg antibody were used per IP.

## Acknowledgement

We thank all members from the Chen and Song labs for the helpful discussion of the project. We thank Xibin Lu for the excellent support of FACS. We acknowledge the assistance of SUSTech Core Research Facilities. The computational work was supported by Center for Computational Science and Engineering at Southern University of Science and Technology.

## Author Contributions

J.Z and X.C conceived the project. J.Z and X.C designed the experiments. J.Z performed the experiment and computational analyses with the help from Y.C, Y.H, J.L, Y.Z and M.X. W.S and X.C supervised the entire project.

## Data Availability

Sequencing data are deposited into ArrayExpress and the accession numbers are E- MTAB-13855 (ChIP-seq), E-MTAB-13856 (bulk RNA-seq) and E-MTAB-13858 (scRNA- seq).

## Code Availability

The code used for the data processing and analysis mentioned in the method section is available in the GitHub repository https://github.com/sibszheng/VZ_development.

---

**Supplementary Figure 1.**
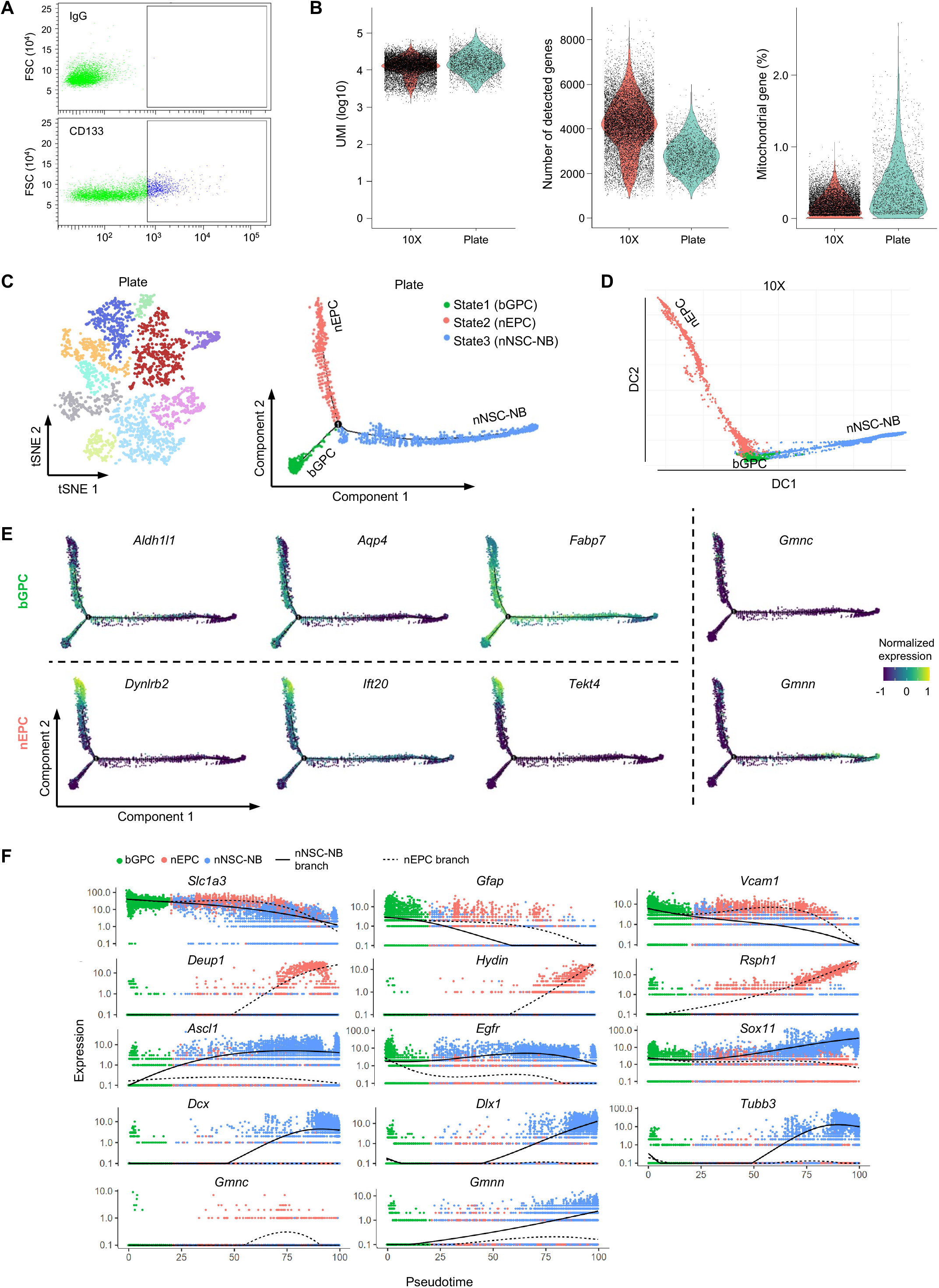
Quality control, trajectory inference and marker gene expression of scRNA-seq. **(A)** Distribution and sorting gates of CD133-labeled cells. IgG served as a negative control for defining the sorting region (indicated by the rectangle in the bottom plot). **(B)** UMI count, gene count, and mitochondrial gene percentage for each individual cell in the 10X and the Plate data. **(C)** 2D t-SNE visualisation (left) of 2,594 cells from the Plate data and the bifurcating trajectory (right) constructed by Monocle. Individual cells are colour-coded according to clusters or states. **(D)** The bifurcating trajectory inferred using the diffusion map algorithm. **(E)** Expression profiles of additional markers used to assign cell classifications of bGPCs, nEPCs and nNSC-NBs. **(F)** Expression dynamics of various marker genes over pseudotime.

**Supplementary Figure 2.**
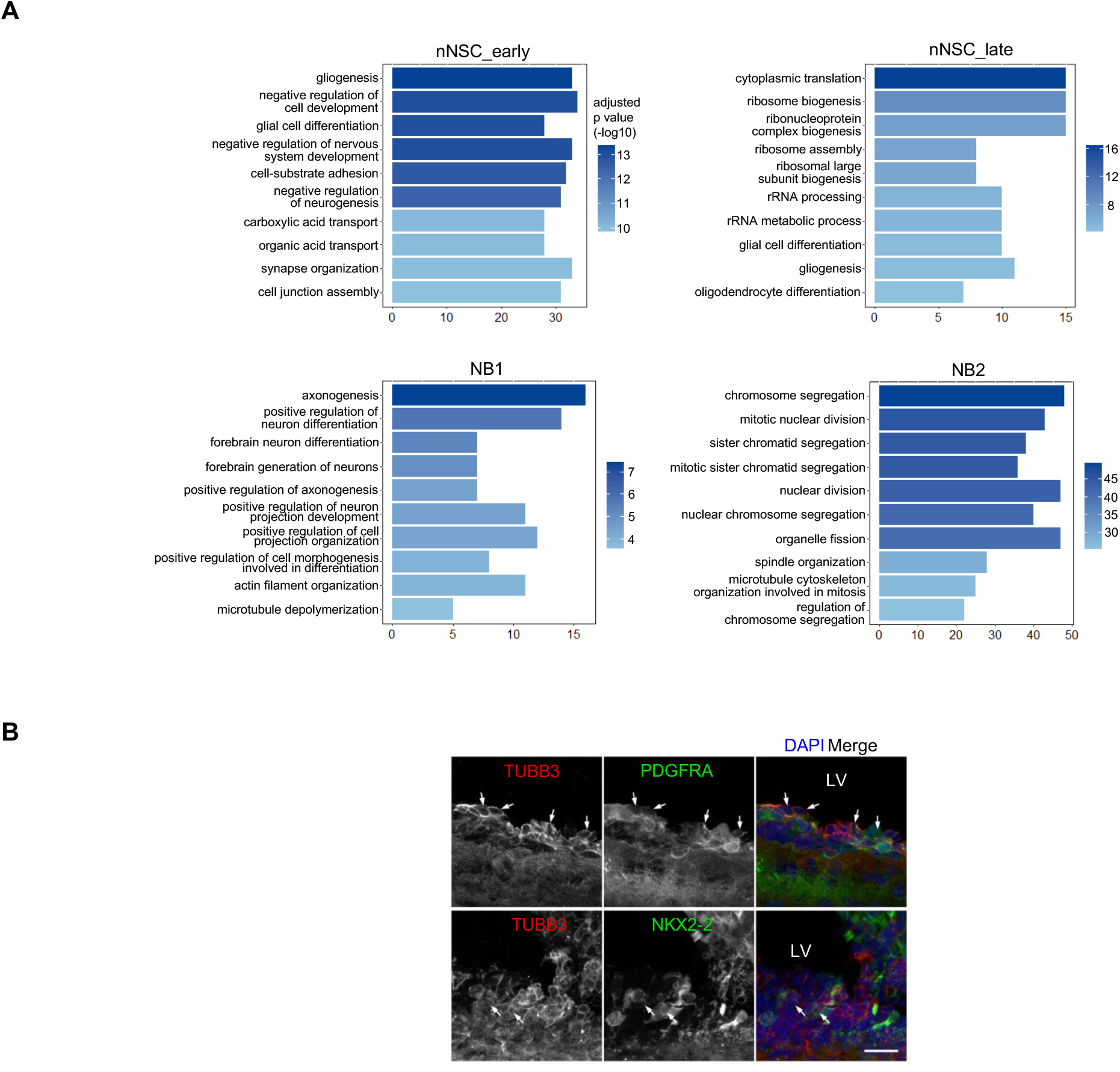
GO analysis of the nNSC-NB branch and the validation of OPC-NB bipotent precursors. **(A)** Top 10 GO terms of biological processes from genes differentially expressed in the four clusters along the nNSC- NB branch. **(B)** Immunofluorescence analyses of neonatal brain sections showing the co-expression of the NB marker (TUBB3) and the OPC marker (PDGFRA or NKX2-2). LV, lateral ventricle. The scale bar is 20 μm.

**Supplementary Figure 3.**
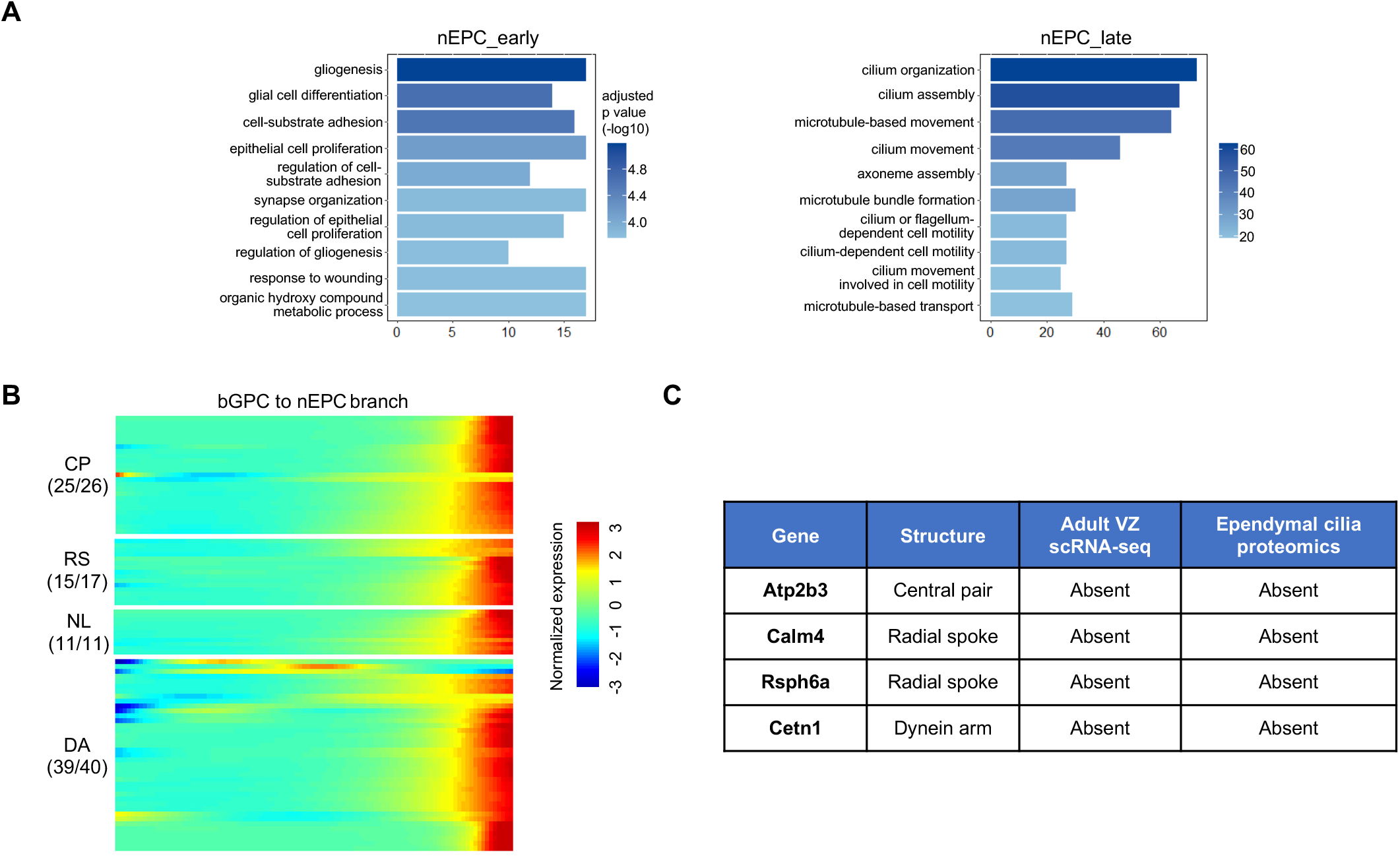
Multiciliogenesis program dominates bGPC transition into nEPCs. **(A)** Top 10 GO terms of biological processes from genes differentially expressed in the two clusters along the nEPC branch. **(B)** Heatmap showing the dynamic expression of multicilia-specific genes along the nEPC branch. **(C)** The four multicilia-specific genes absent in our data were also not detectable in the adult VZ scRNA-seq and ependymal cilia proteomic data.

**Supplementary Figure 4.**
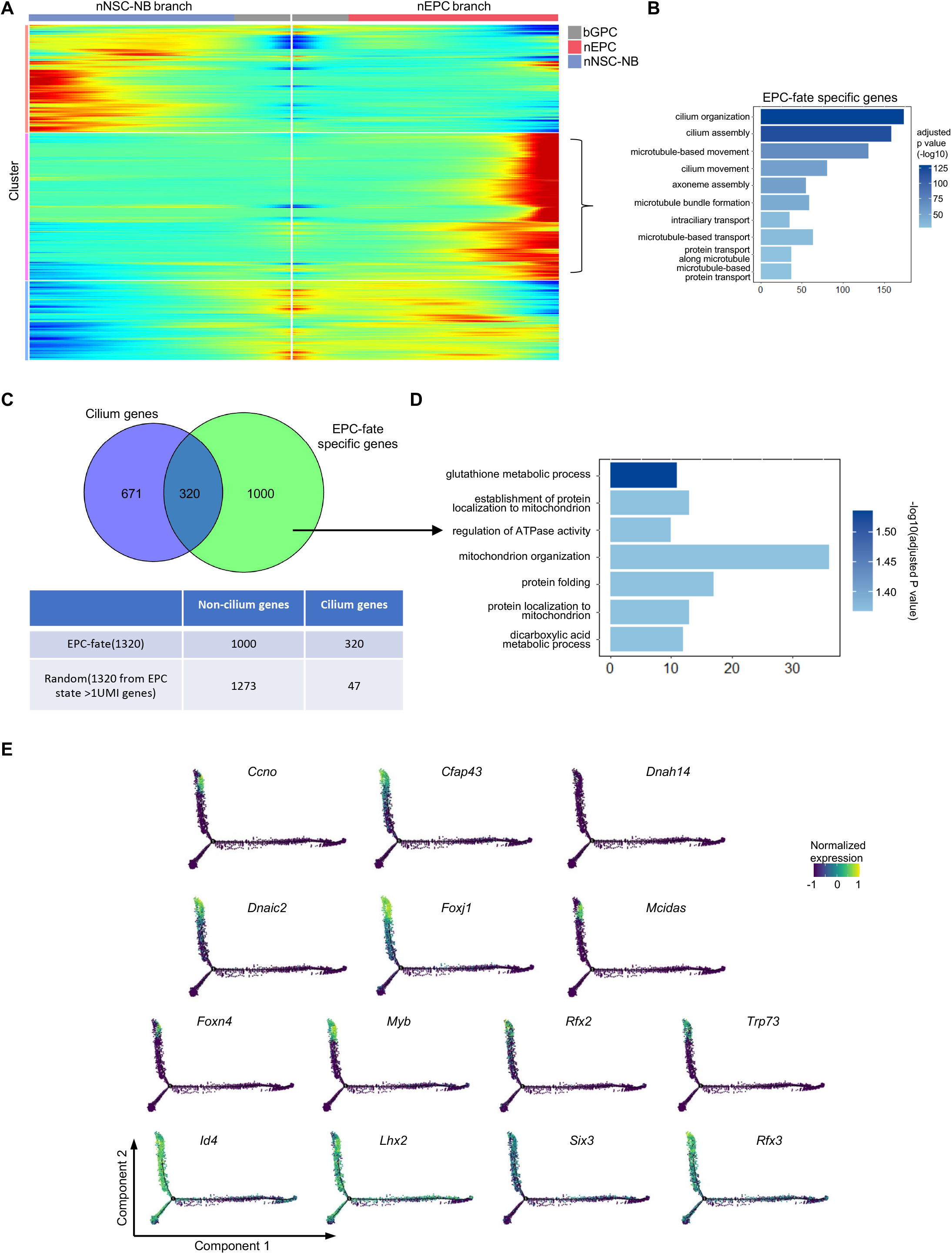
Analysis of EPC-fate specific genes. **(A)** Heatmap showing the dynamic expression of the top 3000 branch-specific differentially expressed genes. **(B)** Top 10 GO terms of biological processes enriched in 1320 EPC-fate specific genes. **(C)** Venn diagram illustrating that 320 EPC-fate specific genes are included in the cilium gene set. **(D)** Top Go terms of biological processes enriched in 1000 non-ciliary EPC-fate specific genes. **(E)** Expression profiles of hydrocephalus genes and EPC-fate regulators in the bifurcating trajectory.

**Supplementary Figure 5.**
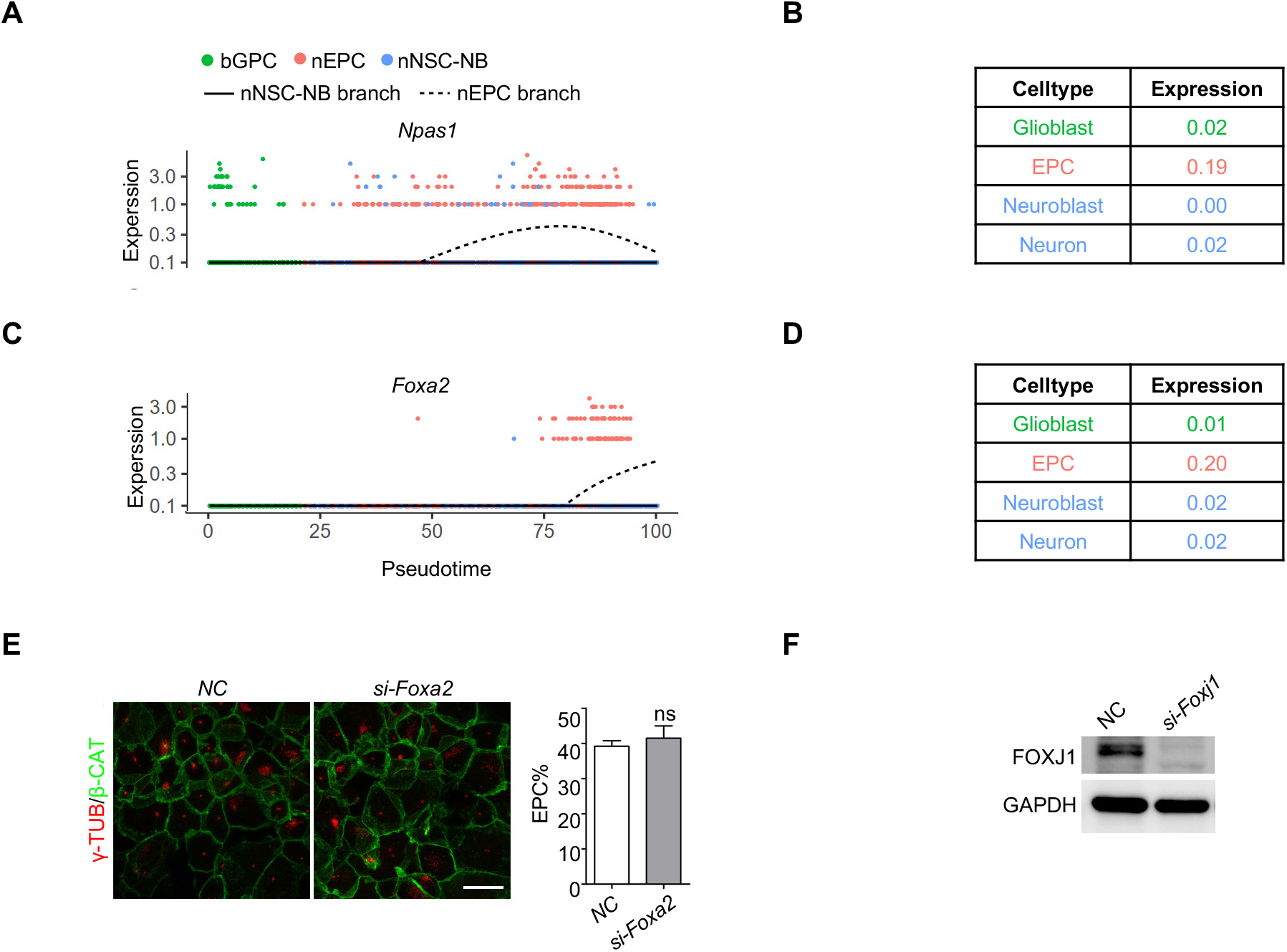
*Npas1* and *Foxa2* are more abundant in EPCs. **(A** and **C)** Expression dynamics of *Npas1* **(A)** and *Foxa2* **(C)** along the pseudotime. **(B** and **D)** Expression values of *Npas1* **(B)** and *Foxa2* **(D)** in the indicated cell types of the developing mouse brain dataset. **(E)** Centriole amplification was not affected by *Foxa2* depletion in EPCs. Error bars represent SD. ns, not significant by the Student’s t-test. The scale bar is 20 μm. **(F)** Immunoblotting showed efficient depletion of *Foxj1* in EPCs by RNAi.

**Supplementary Figure 6.**
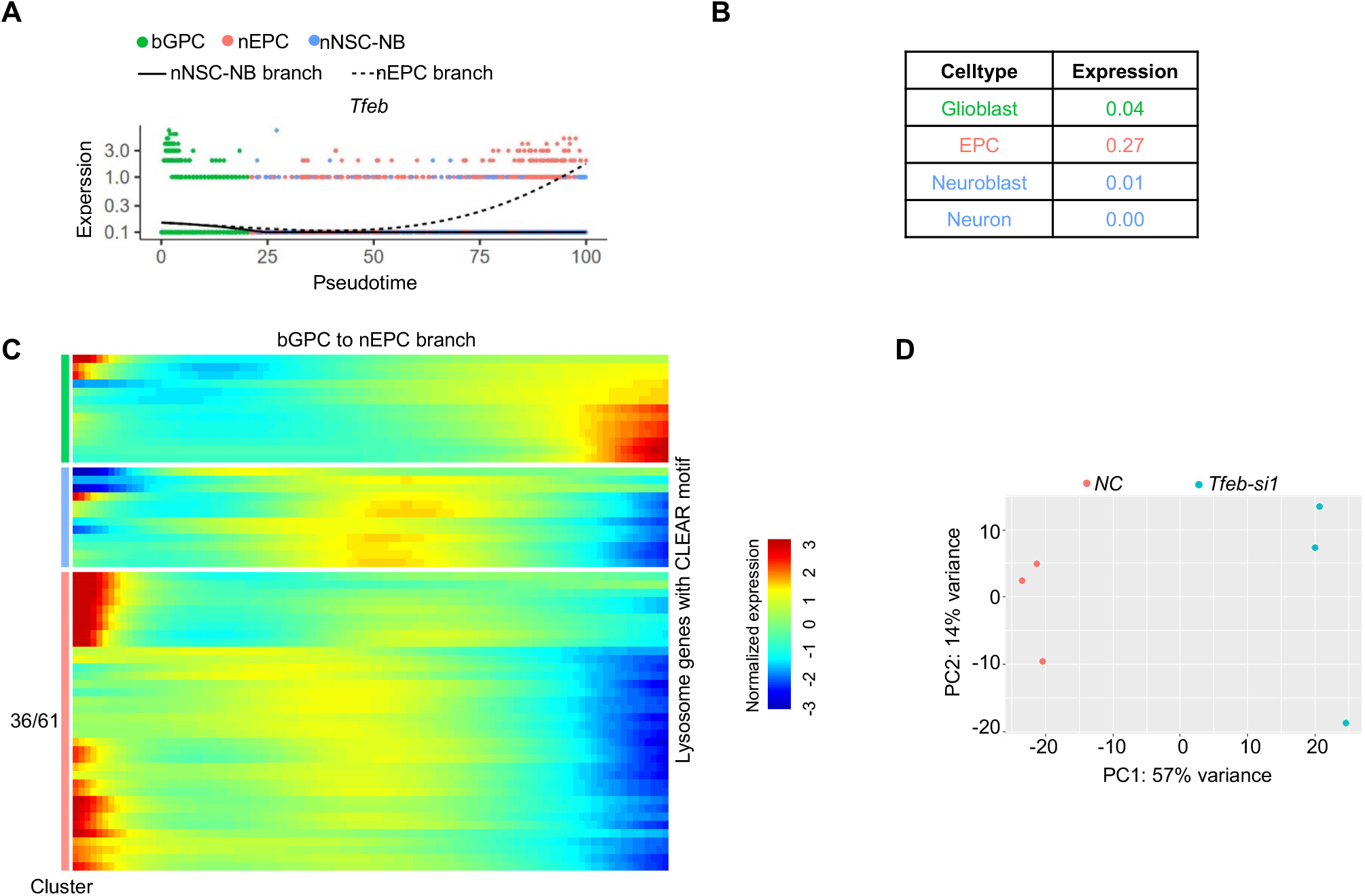
Lysosomal gene expressions during EPC-lineage differentiation are independent of *Tfeb*. **(A)** Expression dynamics of *Tfeb* along the pseudotime. **(B)** Expression values of *Tfeb* in the indicated cell types of the developing mouse brain dataset. **(C)** Heatmap shows the dynamic expression of lysosome genes with CLEAR motif along the nEPC branch. **(D)** Principal component analysis of three replicates of non-targeting control and *Tfeb-si1* samples from the bulk RNA-seq experiments.

**Supplementary Figure 7.**
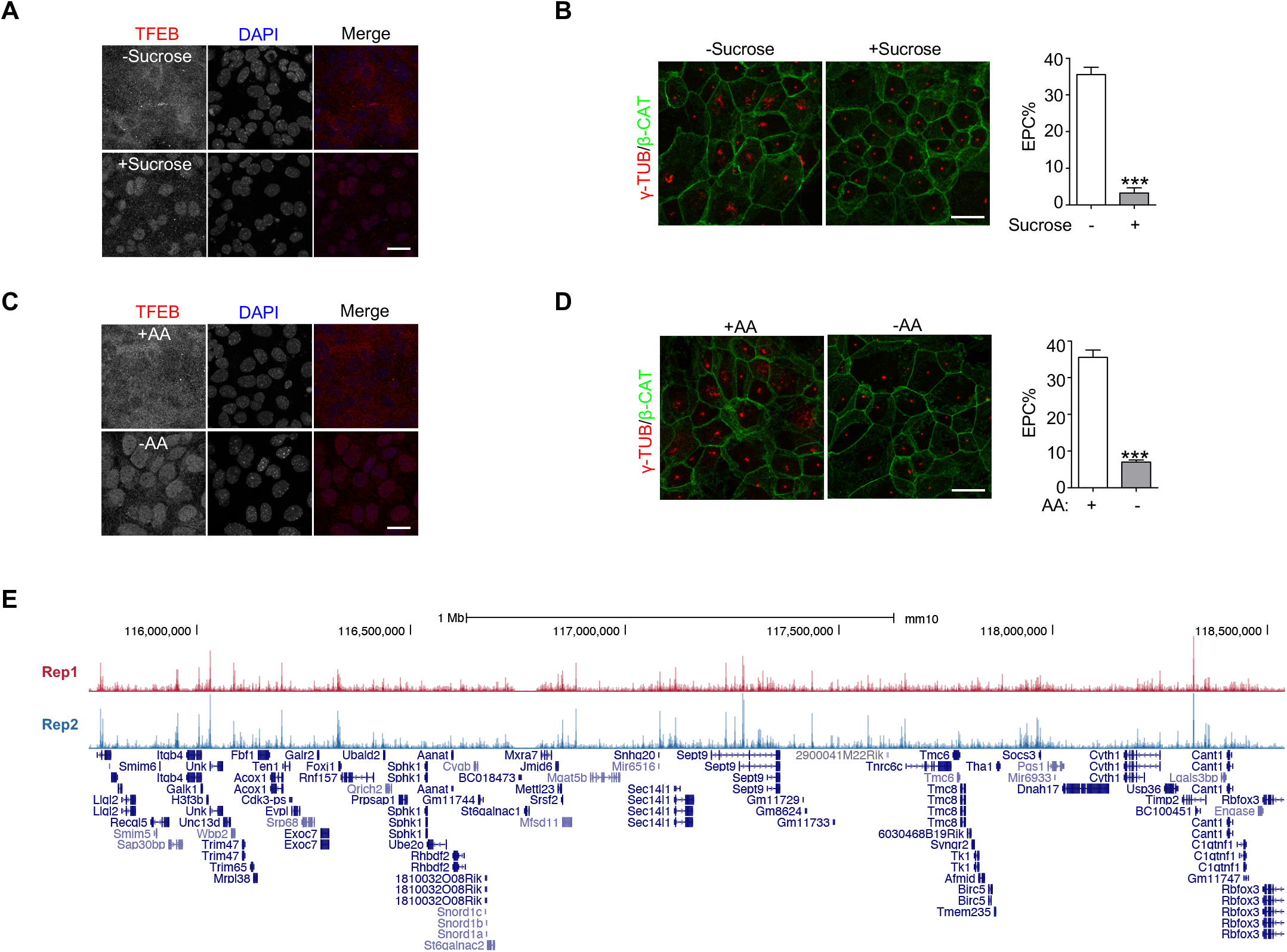
Activation of TFEB blocks the GPC to EPC differentiation. **(A)** TFEB translocated from cytosol to nucleus after 100 mM sucrose treatment. **(B)** A dramatic reduction in the EPC percentage after 100 mM sucrose treatment. **(C)** Amino acid starvation induced translocation of TFEB from the cytosol to the nucleus. **(D)** EPC-lineage differentiation was drastically suppressed after amino acid starvation. Error bars represent the SD. Asterisks indicate P-values from Student’s t-tests, *** P < 0.001. The scale bars are 20 μm. **(E)** UCSC genome browser track showing the good quality of the TFEB ChIP-seq data.

**Supplementary Figure 8.**
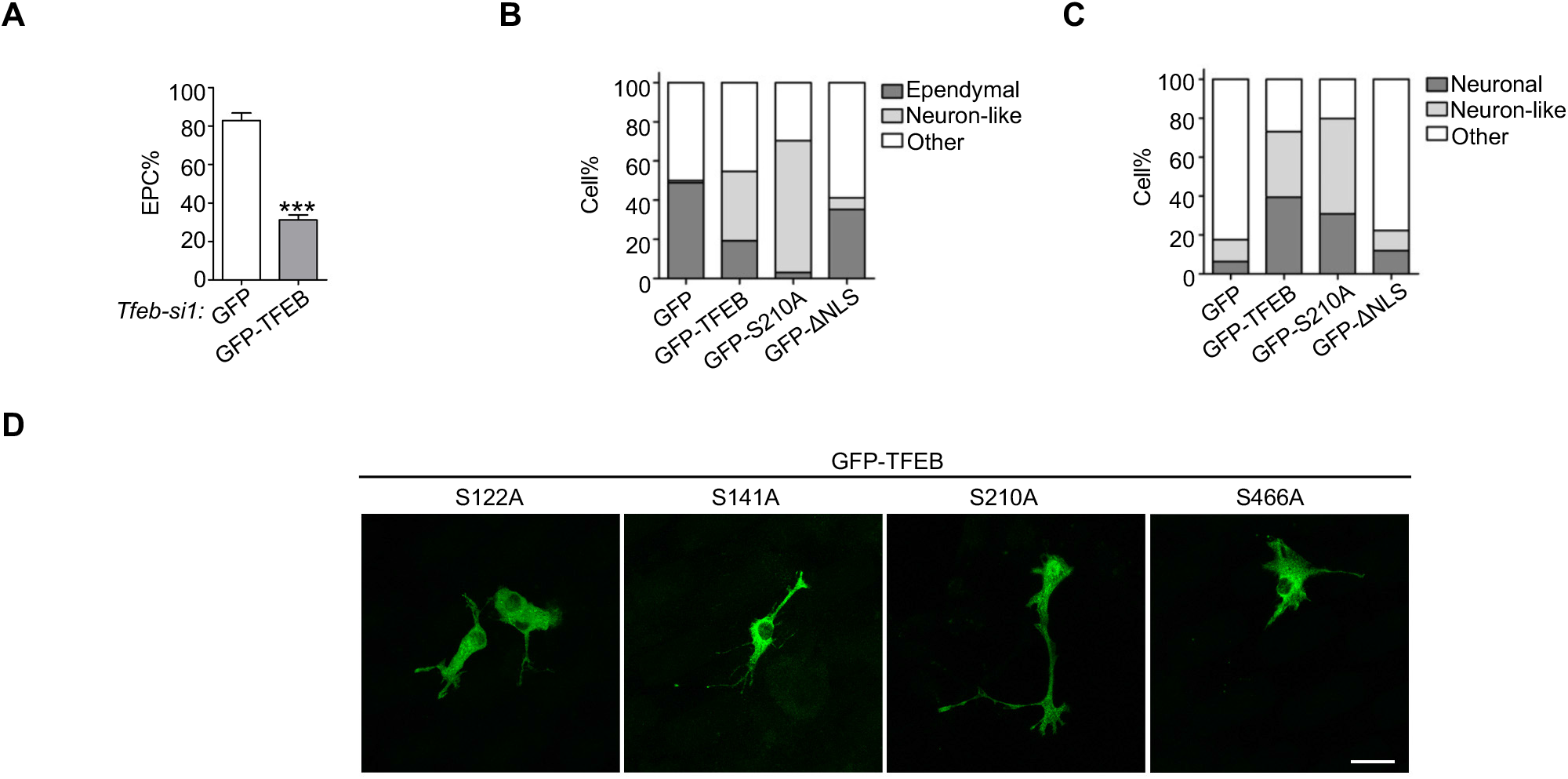
Activation of TFEB induces NSC-lineage differentiation. **(A-C)** Quantifications of Figure 8A-C. Error bars represent SD. Asterisks indicate P-values from Student’s t-test, *** P < 0.001. **(D)** Among the four reported consecutive active mutants of TFEB, only TFEB-S210A showed predominant nuclear localisation when transfected into GPCs. The scale bar is 25 μm.

